# Multiple effectors trigger nonhost resistance in *Solanum americanum* against *Pseudomonas syringae*

**DOI:** 10.1101/2025.05.01.651788

**Authors:** Jieun Kim, Markéta Vlková-Žlebková, Honour C McCann, Kee Hoon Sohn

## Abstract

Wild plant species are threatened by diverse pathogens, but disease symptoms are rarely observed in nature. This suggests that wild plants harbor valuable sources of resistance. In this study, we show that a model bacterial pathogen *Pseudomonas syringae* pv. *tomato* (*Pto*) DC3000 triggered defense responses in all tested accessions of a wild Solanaceae species, *Solanum americanum*. *Pto* DC3000-triggered immunity in *S. americanum* required type III secretion system. We show that seven *Pto* DC3000 effectors (AvrPto, HopAD1, HopAM1, HopC1, HopAA1-1, HopM1, and AvrE1) triggered hypersensitive responses (HR) in *S. americanum* accession SP2273. Significantly, sequential deletion of the HR-triggering effectors from *Pto* DC3000 resulted in enhanced virulence in *S. americanum*. However, the well-conserved effectors, HopM1 and AvrE1 were indispensable for virulence. We conclude that the immunity triggered by multiple effectors contributes to nonhost resistance in *S. americanum* against *P. syringae*. We propose that the identification of the corresponding disease resistance genes for HopM1 and AvrE1 in *S. americanum* would accelerate development of durable immunity to *P. syringae* pathogens in Solanaceae crops.

## INTRODUCTION

Plants are constantly threatened by the invasion of pathogens, yet plant diseases are relatively uncommon in nature (Gill et al., 2015). This is due to the diverse defense strategies that plants have evolved. There are two major immune layers (Jones and Dangl, 2006). The first layer is pattern-triggered immunity (PTI), where pattern recognition receptors (PRRs) localized at the plant cell surface detect conserved pathogen-associated molecules such as bacterial flagellin. However, bacterial pathogens have developed sophisticated strategies to evade PTI by delivering effector proteins to plant cells via type III secretion system (Macho, 2016). Many bacterial effectors contribute to pathogen virulence by suppressing basal plant immunity. For example, a well-studied *Pseudomonas syringae* effector, AvrPto interferes the kinase function of certain PRRs such as *Arabidopsis* FLS2 or EFR, resulting in reduced PTI signaling (Xiang et al., 2008; Zipfel and Rathjen, 2008). In response to these effector activities, plants have evolved a second layer of innate immunity termed effector-triggered immunity (ETI) where nucleotide-binding and leucine-rich repeat resistance (NLR) proteins recognize corresponding pathogen effectors (Jones and Dangl, 2006; Kourelis and van der Hoorn, 2018). ETI often leads to hypersensitive response (HR), characterized by localized cell death at the infection site. Some NLR proteins, such as HOP<u>Z</u>-<u>A</u>CTIVATED <u>R</u>ESISTANCE<u>1</u> (ZAR1), oligomerize upon effector recognition and function as a calcium-permeable channel in the plasma membrane which is required for HR development (Bi et al., 2021; Wang et al., 2019).

*Pseudomonas syringae* pv. *tomato* DC3000 (hereafter *Pto* DC3000) is a model bacterial pathogen for studying plant-microbe interactions (Buell et al., 2003; Lindeberg et al., 2006; Xin and He, 2013). In particular, the secretion and *in planta* functions of *Pto* DC3000 type III effectors have been extensively studied. *Pto* DC3000 secretes multiple effectors whose functions can be complex and redundant. This nature makes it challenging to study individual effector functions. To overcome these difficulties, combinations of multiple effectors were deleted in *Pto* DC3000 background (Wei et al., 2007). Among 36 known *Pto* DC3000 type III effectors, 28 are well-expressed (18 of these are clustered in six loci, while 10 effectors are dispersed throughout the genome). The remaining eight effectors are either pseudogenes or weakly expressed genes. Deletion of 18 clustered and well-expressed effector genes in *Pto* DC3000 resulted in D18E, and deletion of all 28 well-expressed effector genes produced D28E (Cunnac et al., 2011; Kvitko et al., 2009). These *Pto* DC3000 polymutant strains showed significantly reduced virulence in a model Solanaceae species *Nicotiana benthamiana*. Further deletion of a weakly-expressed effector *hopAD1* in D28E, resulted in D29E, which abolished HR in *N. benthamiana* (Wei et al., 2015). Finally, the effectorless mutant D36E was generated by deleting all the remaining weakly expressed effectors and pseudogenes from D29E (Wei et al., 2015). *Pto* DC3000 effectors encoded by Exchangeable Effector Locus (EEL) vary among different strains, whereas Conserved Effector Locus (CEL) encodes highly conserved effectors such as AvrE1, HopM1, HopAA1 (HopN1 in some strains) across diverse *P. syringae* strains (Alfano et al., 2000; Xin et al., 2018). Since CEL effectors are highly conserved, identifying their corresponding resistance genes is considered to be crucial for developing durable disease resistance to *P. syringae* pathogens (Dangl et al., 2013; Kim et al., 2022).

Wild plant species are valuable sources of resistance genes compared to domesticated crops (Arora et al., 2019). Several *P. syringae* effectors have been shown to activate NLR-mediated immunity in *N. benthamiana*, a model Solanaceae species. HopQ1 effector from *P. syringae*, for example, is recognized by NLR protein Roq1 with N-terminal Toll-like interleukin-1 (TIR) domain (Schultink et al., 2017). Interestingly, Roq1 also recognizes *Xanthamonas* and *Ralstonia* effectors, XopQ and RipB, respectively. Recently, *N. benthamiana* and *Solanum lycopersicoides* NLR Ptr1, which contains coiled-coil (CC) domain, was shown to recognize multiple bacterial effectors. For instance, *P. syringae* effectors AvrRpt2, AvrRpm1, AvrB, and HopZ5 activate NbPtr1-dependent immunity in *N. benthamiana* (Ahn et al., 2023). Moreover, NbPtr1 also recognizes RipBN and RipE1 from *Ralstonia solanacearum,* and AvrBsT from *Xanthomonas euvesicatoria*. *Solanum americanum*, a wild Solanaceae species, has been used to identify NLR genes (Witek et al., 2016). High-quality genome assemblies and NLR gene repertoires have been analyzed in multiple *S. americanum* accessions in recent research (Lin et al., 2023; Witek *et al*., 2016). Moreover, the availability of genetically variable *S. americanum* accessions makes *S. americanum* an ideal model for discovering NLR genes (Witek *et al*., 2016). Several NLR genes that recognize effectors from *Phytophthora infestans* causing potato late blight have been identified and cloned from *S. americanum* (Lin *et al*., 2023; Witek *et al*., 2016; Witek et al., 2021). For instance, Rpi-amr1 from *S. americanum* recognizes Avr-amr1 and provides resistance when expressed in other Solanaceae species, such as potato and *N. benthamiana*. Additionally, Rpi-amr3 recognizing Avr-amr3 confers broad resistance to *P. infestans* and other *Phytophthora* pathogens including *P. parasitica* and *P. palmivora* when expressed in *N. benthamiana* (Lin et al., 2022).

In this study, we aimed to understand the genetic basis of *S. americanum* resistance to *Pto* DC3000 and found that seven effectors (AvrPto, HopAD1, HopAM1, HopC1, HopAA1-1, HopM1, and AvrE1) trigger HR in *S. americanum* accession SP2273. We generated *Pto* DC3000 mutants lacking these HR-triggering effectors and investigated their roles in bacterial disease resistance *in S. americanum*. We show that the deletion of *avrPto*, *hopAD1*, *hopAM1*, *hopC1*, and *hopAA1-1* from *Pto* DC3000 enhances *in planta* bacterial growth and causes bacterial speck symptoms while still inducing weak HR in *S. americanum*. Deletion of all seven avirulence effectors abolished the HR-triggering ability of *Pto* DC3000 in *S. americanum* without disease development. Based on these results, we propose that multiple effectors are required for nonhost resistance in *S. americanum* against *P. syringae*. Our results offer insights into bacterial virulence mechanisms and can help the identification of NLRs recognizing CEL effectors, which may confer durable resistance in Solanaceae crops.

## RESULTS

### *Pseudomonas syringae* pv. *tomato* DC3000 triggers type III secretion system-dependent disease resistance in *Solanum americanum*

To better understand the genetic basis of disease resistance to *Pseudomonas syringae* in *S. americanum*, we infiltrated *Pseudomonas syringae* pv. *tomato* (*Pto*) DC3000 (OD_600nm_=0.1) into the leaves of 28 *S. americanum* accessions and tested for the onset of hypersensitive response (HR). Unlike *P. infestans* and *R. pseudosolanacearum,* which induce accession-specific resistance in *S. americanum* (Moon et al., 2021; Witek *et al*., 2016), *Pto* DC3000 wild-type strain triggered a strong HR in all tested *S. americanum* accessions at one day post-infection (dpi) (Table 1). We hypothesized that one or more *Pto* DC3000 type III effector (T3E) proteins trigger HR in *S. americanum*. In order to identify which *Pto* DC3000 T3E(s) are responsible for HR, we tested HR-inducing activity of *Pto* DC3000 polymutants D18E, D29E, and D36E lacking multiple T3Es (Kvitko *et al*., 2009; Wei *et al*., 2015) in *S. americanum* accession SP2273 (hereafter, SP2273). *Pto* DC3000 wild-type and D18E triggered strong HR, whereas D29E and D36E did not induce any visible symptoms in SP2273 (Figure 1A). Next, we tested the virulence of *Pto* DC3000 wild-type and polymutants by measuring *in planta* bacterial growth. None of the strains (*Pto* DC3000 wild-type, D18E, D29E, or D36E) showed growth in SP2273 (Figure 1B), suggesting that deletion of multiple T3Es resulted in the loss of not only HR but also bacterial virulence in *S. americanum*. Based on these results, we hypothesized that the 11 effectors present in D18E but absent in D29E (HopA1, HopAD1, HopAF1, HopAM1, HopB1, HopE1, AvrPto, AvrPtoB, HopI1, HopK1, and HopY1) are primary avirulence effector candidates (Figure 1C). In addition, the 18 effectors that are present in wild-type strain but absent in D18E (HopAA1-1, HopAA1-2, HopAO1, AvrE1, HopC1, HopD1, HopR1, HopG1, HopH1, HopM1, HopN1, HopO1-1, HopQ1-1, HopF2, HopT1-1, HopU1, HopV1, and HopX1) could be the additional avirulence effector candidates (Fig 1C). We excluded the seven effectors (HopS2, HopO1-3’, HopO1-2, HopS1’, HopBM1, HopT2, and HopT1-2’) remaining in D29E because D29E failed to trigger HR, and these genes are considered weakly expressed genes or pseudogenes (Wei *et al*., 2015). Taken together, these results strongly suggest that T3Es are essential for *Pto* DC3000-induced HR in *S. americanum*.

**Figure 1.**
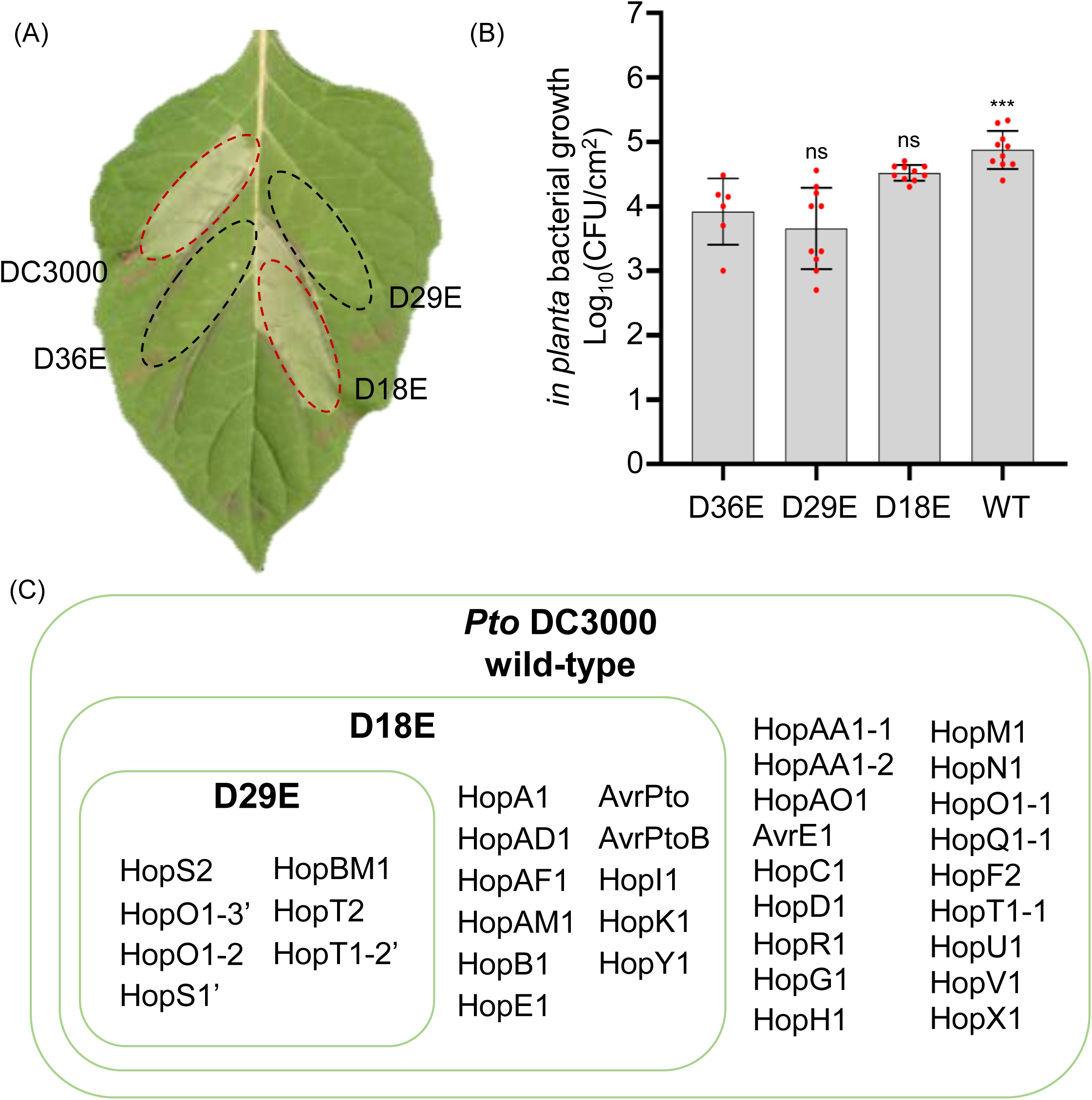
*Pseudomonas syringae* pv. *tomato* DC3000 polymutants show type III effector-dependent phenotypes in *Solanum americanum*. **(A)** HR phenotypes triggered by *Pto* DC3000 wild-type and polymutants. Strains were infiltrated using a needleless syringe (OD_600nm_=0.1) into *S. americanum* accession SP2273 leaves. Photographs were taken one day post infiltration. Red dashed borders indicate HR, and black dashed borders indicate no HR within the infiltrated area. **(B)** *In planta* bacterial growth of *Pto* DC3000 wild-type and polymutants in *S. americanum*. Bacteria were infiltrated using a needleless syringe at OD_600nm_=0.0001 in SP2273. *In planta* bacterial growth was quantified four days post infiltration. Individual data points are represented by red dots, and the bars indicate standard deviation. Raw data is presented in Table S4. Statistical significance was determined using one-way ANOVA followed by Dunnett’s multiple tests (ns: nonsignificant, **: p<0.01, ***: p<0.001). GRAPHPAD PRISM v.10.0.1 was used for statistical analysis. **(C)** Type III effector repertoires in *Pto* DC3000 polymutants. Effectors within the box are the remaining effectors in each *Pto* DC3000 polymutant.

### AvrPto, HopAM1, or HopAD1 triggers hypersensitive response in *Solanum americanum*

To identify avirulence effector(s) that trigger defense responses in *S. americanum*, we transiently expressed each of the 11 effectors present in D18E (HopA1, HopAD1, HopAF1, HopAM1, HopB1, HopE1, AvrPto, AvrPtoB, HopI1, HopK1, and HopY1) in SP2273 leaves by *Agrobacterium*-mediated transient transformation (hereafter, agroinfiltration). Among these, agroinfiltration of HopAD1, HopAM1, or AvrPto elicited a robust HR in SP2273 (Figure 2A). To test whether the lack of HR for other effectors was due to protein instability. We transiently expressed all 11 effectors in *Nicotiana benthamiana* leaves via agroinfiltration and total protein extracts were analyzed by immunoblot using anti-HA antibody. All tested effectors showed detectable protein expression levels (Figure 2B and Table S5). Although HopAM1 showed weaker expression compared to other effectors, it still induced HR (Figure 2A), indicating that the level of expression was sufficient to activate plant defense responses.

**Figure 2.**
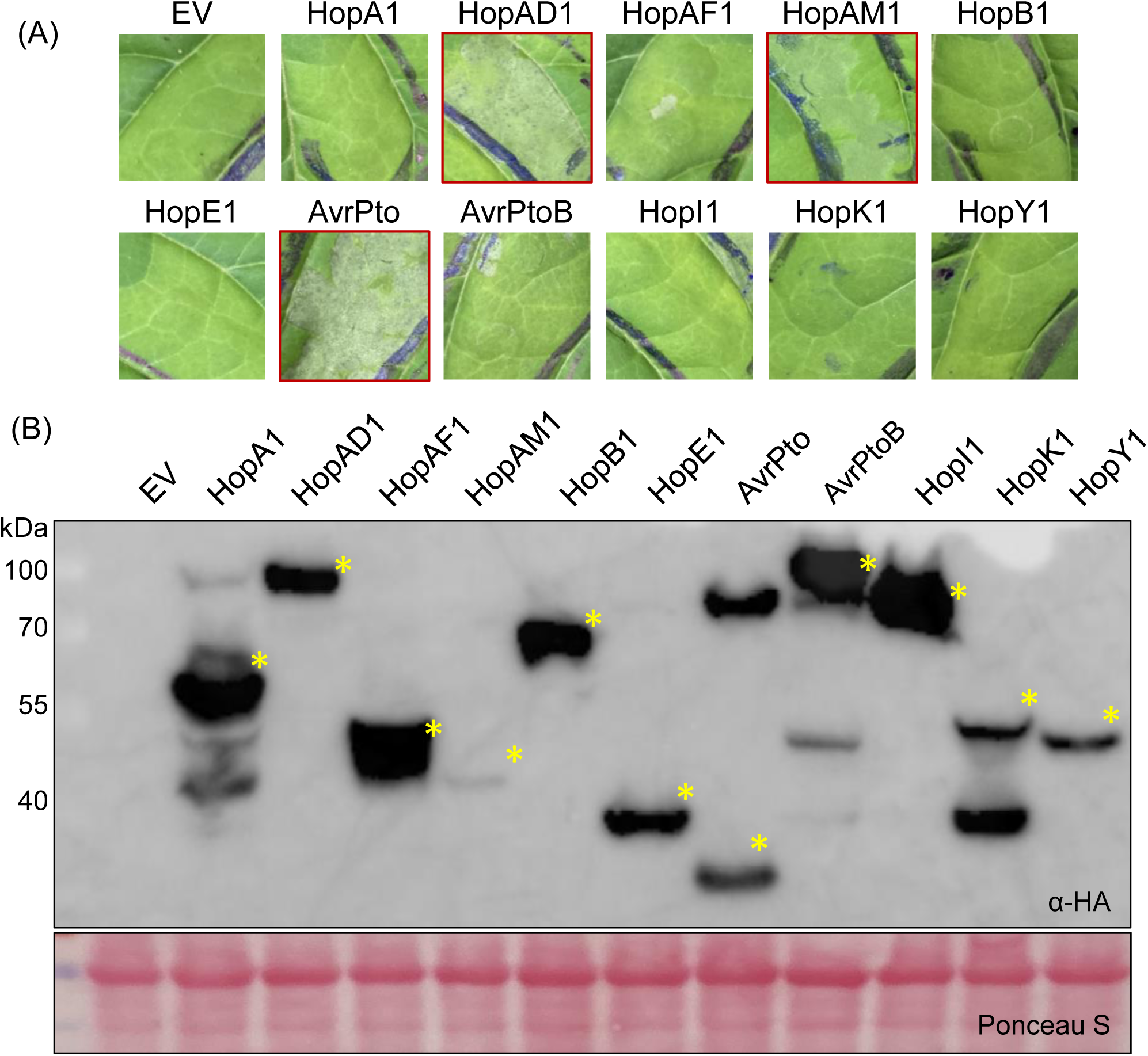
AvrPto, HopAM1, or HopAD1 triggers a hypersensitive response in *Solanum americanum.* **(A)** 11 effectors present in D18E but not in D29E were transiently expressed in SP2273. *Agrobacterium* carrying C-terminal HA-tagged effectors was infiltrated into SP2273 at OD_600nm_=0.4 with P19 (OD_600nm_=0.2), which is suppressor of gene silencing. Photographs were taken four days post infiltration. Red borders indicate the presence of HR. **(B)** Protein accumulation of effectors was confirmed by immunoblot assay. *Agrobacterium* carrying C-terminal HA-tagged type III effectors and P19 were co-infiltrated into *Nicotiana benthamiana* leaves. The inoculum concentration of the effector was OD_600nm_=0.4 and P19 concentration was OD_600nm_ = 0.2. Leaf samples were collected two days post infiltration. Protein accumulation was detected using an anti-HA antibody. Yellow asterisks indicate expected protein bands. Ponceau S staining shows an equal amount of protein loading.

### *Pto* DC3000 lacking *avrPto*, *hopAM1*, and *hopAD1* shows enhanced *in planta* bacterial growth, yet still triggers hypersensitive response in *Solanum americanum*

To investigate the roles of AvrPto, HopAM1, and HopAD1 in *Pto* DC3000 virulence, we sequentially deleted each effector gene and tested for HR induction and *in planta* bacterial growth in *S. amerianum*. Effector knock-out mutants were generated using a modified suicide vector pK18mobsacB-GG containing the upstream and downstream flanking regions of the target effector genes (Jayaraman et al., 2020). First, we deleted *avrPto* from *Pto* DC3000 wild-type resulting in strain PKSG 4673 (Figure 3A). Since *hopAM1* gene exists in two identical copies, one on the chromosome (*hopAM1-1*) and the other on the plasmid, pDC3000A (*hopAM1-2*) (Buell *et al*., 2003), we deleted *hopAM1-1* from PKSG 4673 (PKSG 7065) and subsequently deleted *hopAM1-2* from PKSG 7065 (PKSG 7903). Finally, we deleted *hopAD1* from PKSG 7903, generating PKSG 7377. Details on the generation and validation of these effector knockout strains are provided in the Methods section and Figure S1.

**Figure 3.**
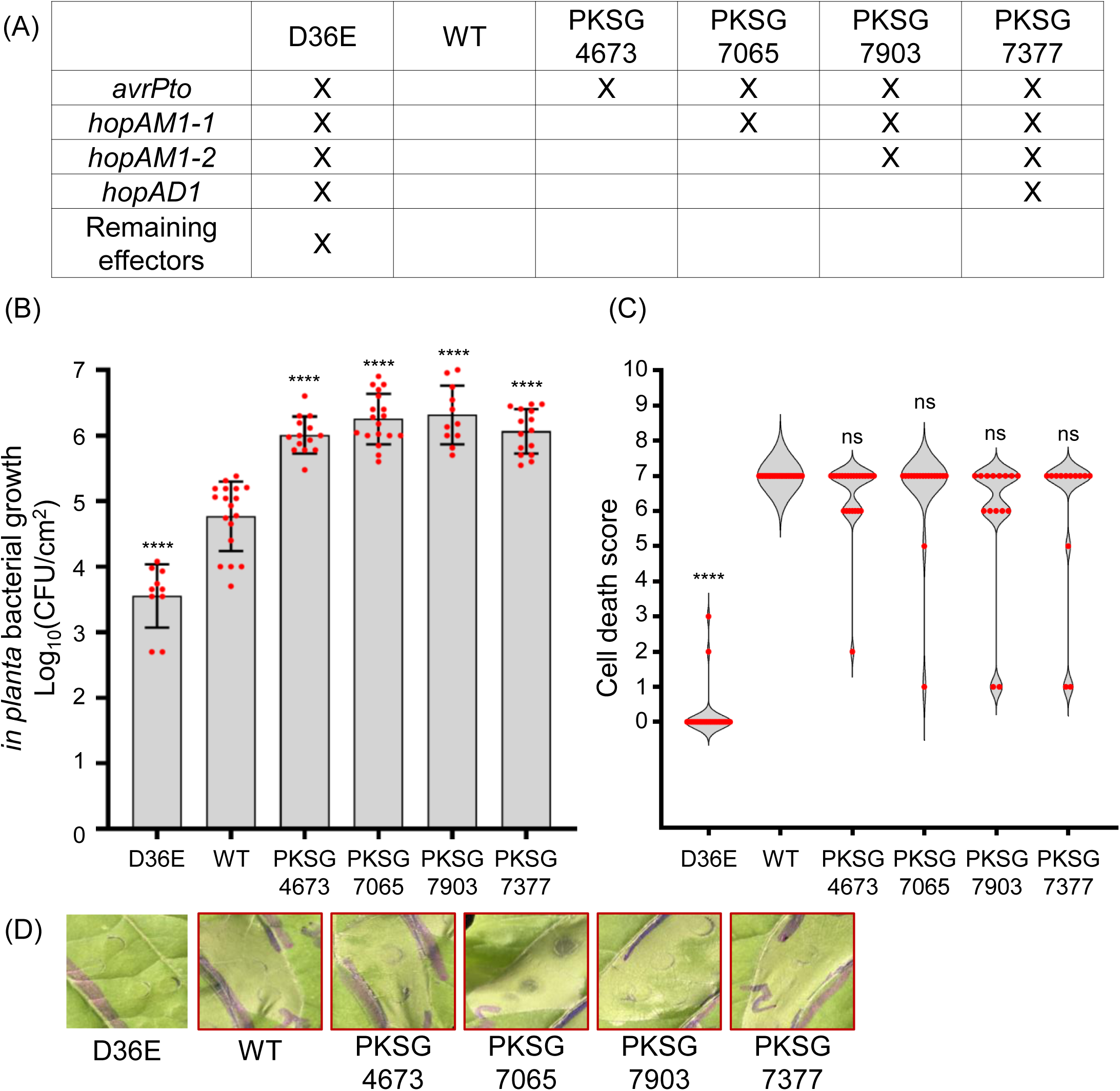
Deletion of avirulence effectors enhances *in planta* bacterial growth but maintains hypersensitive response induction. **(A)** Names of *Pto* DC3000 mutants and deleted effectors. Effector deletions were performed sequentially, as detailed in the Methods section. **(B)** *Pto* DC3000 mutants deleted with HR-triggering effectors show enhanced growth compared to wild-type. Bacterial strains were infiltrated at OD_600nm_=0.0001 using a needleless syringe into SP2273. *In planta* bacterial growth was assessed 4 days post infiltration. Each red dot represents the individual replicate, with the bars indicating the standard deviation. Raw data are shown in Table S6. Statistical significance was determined using one-way ANOVA followed by Dunnett’s multiple tests (ns: nonsignificant, **: p<0.01, ****: p<0.0001). GRAPHPAD PRISM v.10.0.1 was used for statistical analysis. **(C)** *Pto* DC3000 mutants still trigger HR. Bacterial strains were infiltrated at OD_600nm_=0.1 using a needleless syringe into SP2273 leaves. HR was scored one day post infiltration on a scale from 0 (no HR) to 7 (full HR), represented as violin plots. HR scoring criteria are shown in Figure S2, revised from the previous study (Ahn *et al*., 2023). Individual data points are shown as red dots. Statistical significance was conducted using the Kruskal-Wallis test followed by Dunn’s multiple comparison test (ns: nonsignificant, **: p<0.01, ****: p<0.0001), using GRAPHPAD PRISM v.10.01. Raw data of HR scoring are shown in Table S7. **(D)** Representations of HR phenotypes. Bacterial strains were syringe-infiltrated at OD_600nm_=0.1 into SP2273 leaves. The HR photos were taken one day post infiltration. The red border indicates the presence of HR.

To test whether these effector knockout mutants exhibited enhanced virulence, we measured *in planta* bacterial growth by infiltrating a low concentration of bacterial suspension (OD_600nm_=0.0001) into SP2273 leaves using a needleless syringe. All mutant strains, PKSG 4673 (Δ*avrPto*), PKSG 7065 (Δ*avrPto hopAM1-1*), PKSG 7903 (Δ*avrPto hopAM1-1 hopAM1-2*), PKSG 7377 (Δ*avrPto hopAM1-1 hopAM1-2 hopAD1*) showed significantly higher *in planta* growth compared to wild-type (Figure 3B). However, no significant differences were observed among the mutant strains. To further characterize these mutant strains, we tested their HR-inducing ability by infiltrating a high concentration of bacterial inoculum (OD_600nm_=0.1) into SP2273 leaves. Interestingly, all four *Pto* DC3000 mutant strains still triggered a strong HR in SP2273 (Figure 3C and 3D). Taken together, these results indicate that AvrPto, HopAM1, and HopAD1 significantly contribute to the avirulence of *Pto* DC3000 in *S. americanum.* However, the remaining HR suggests that additional avirulence determinant(s) are likely present in *Pto* DC3000.

### Transient expression of HopC1, HopAA1-1, HopM1 or AvrE1 triggers hypersensitive response in *Solanum americanum*

To identify additional avirulence effector(s), we tested the HR-inducing activity of 18 secondary candidates (HopAA1-1, HopAA1-2, HopAO1, AvrE1, HopC1, HopD1, HopR1, HopG1, HopH1, HopM1, HopN1, HopO1-1, HopQ1-1, HopF2, HopT1-1, HopU1, HopV1, and HopX1) in SP2273 (Figure 1C). Agroinfiltration of these effectors into SP2273 leaves showed that HopAA1-1, AvrE1, HopC1, or HopM1 induced a strong HR at 3 dpi (Figure 4A). HopAA1-1, AvrE1, and HopM1 are known to be highly conserved effectors among diverse *P. syringae* strains (Alfano *et al*., 2000; Munkvold et al., 2009). We next assessed protein expression of these effectors by immunoblot analysis using an anti-HA antibody. Most effectors showed detectable levels of expression. However, for unknown reasons, the expected size bands for AvrE1 (202 kDa) and HopR1 (217 kDa) (Table S5) were not detected in our experimental conditions (Figure 4B).

**Figure 4.**
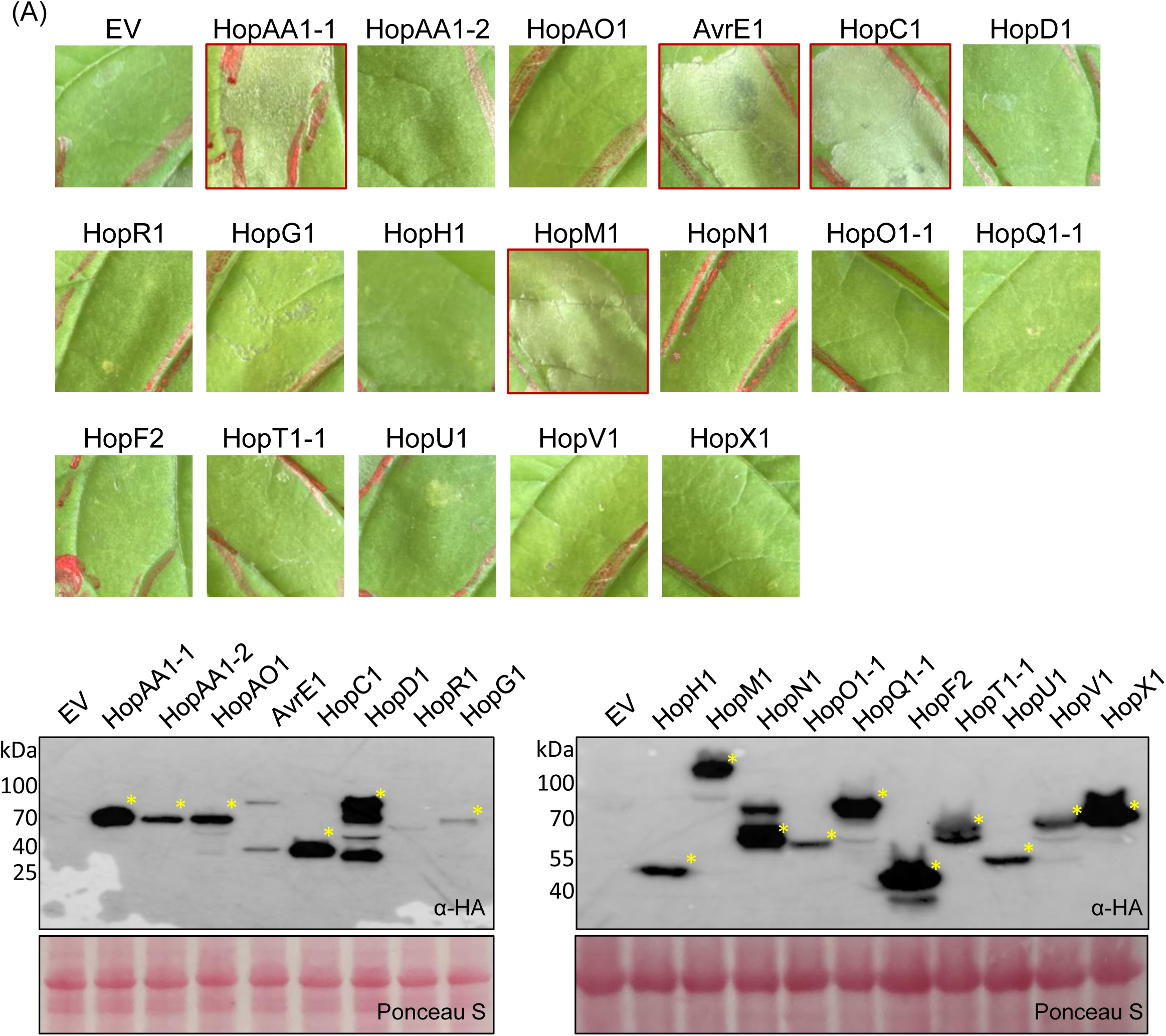
HopAA1-1, AvrE1, HopC1, or HopM1 triggers hypersensitive response in *Solanum americanum.* **(A)** 18 effectors were transiently expressed in *S. americanum* leaves. *Agrobacterium* carrying HA-tagged type III effector and P19 constructs which is suppressor of gene silencing were co-infiltrated into SP2273 at OD_600nm_=0.4 and 0.2, respectively. Photographs were taken four days post infiltration. **(B)** Protein accumulation of effectors was confirmed by immunoblot assay. *Agrobacterium* carrying C-terminal HA-tagged type III effectors and P19 were co-infiltrated in *Nicotiana benthamiana* leaves. The inoculum concentration of the effector was OD_600nm_=0.4, and P19 concentration was OD_600nm_ = 0.2. Leaf samples were collected two days post infiltration. Protein accumulation was visualized using an HA antibody. Yellow asterisks indicate the expected protein size bands. Ponceau S staining shows an equal amount of protein loading.

### HopM1 and AvrE1 are critical for bacterial virulence and disease symptom development in *Solanum americanum*

To test the roles of HopC1, HopAA1-1, HopM1, and AvrE1 in HR induction and bacterial virulence, we generated additional effector knockout mutant strains of *Pto* DC3000. Details for generation and validation of these effector knockout strains can be found in the Methods section and Figure S3. HopM1 and AvrE1 require chaperone proteins ShcM and ShcE, respectively, for proper functions (Badel et al., 2003; Badel et al., 2006). Therefore, *shcM* and *shcE* were deleted along with their corresponding effector genes, *hopM1* and *avrE1*. First, *hopC1* was deleted in PKSG 7377 which resulted in Δ*avrPto hopAM1-1 hopAM1-2 hopAD1 hopC1* (PKSG 7768) (Figure 5A). Next, *hopAA1-1* was deleted in PKSG 7768 to generate PKSG 7826. We further deleted *shcM-hopM1 or/and schE-avrE1* resulting in three additional knockout strains (PKSG 7899; *shcM*-*hopM1* deletion, PKSG 7900; *shcE*-*avrE1* deletion, and PKSG 7892; both deletions) (Figure 5A).

**Figure 5.**
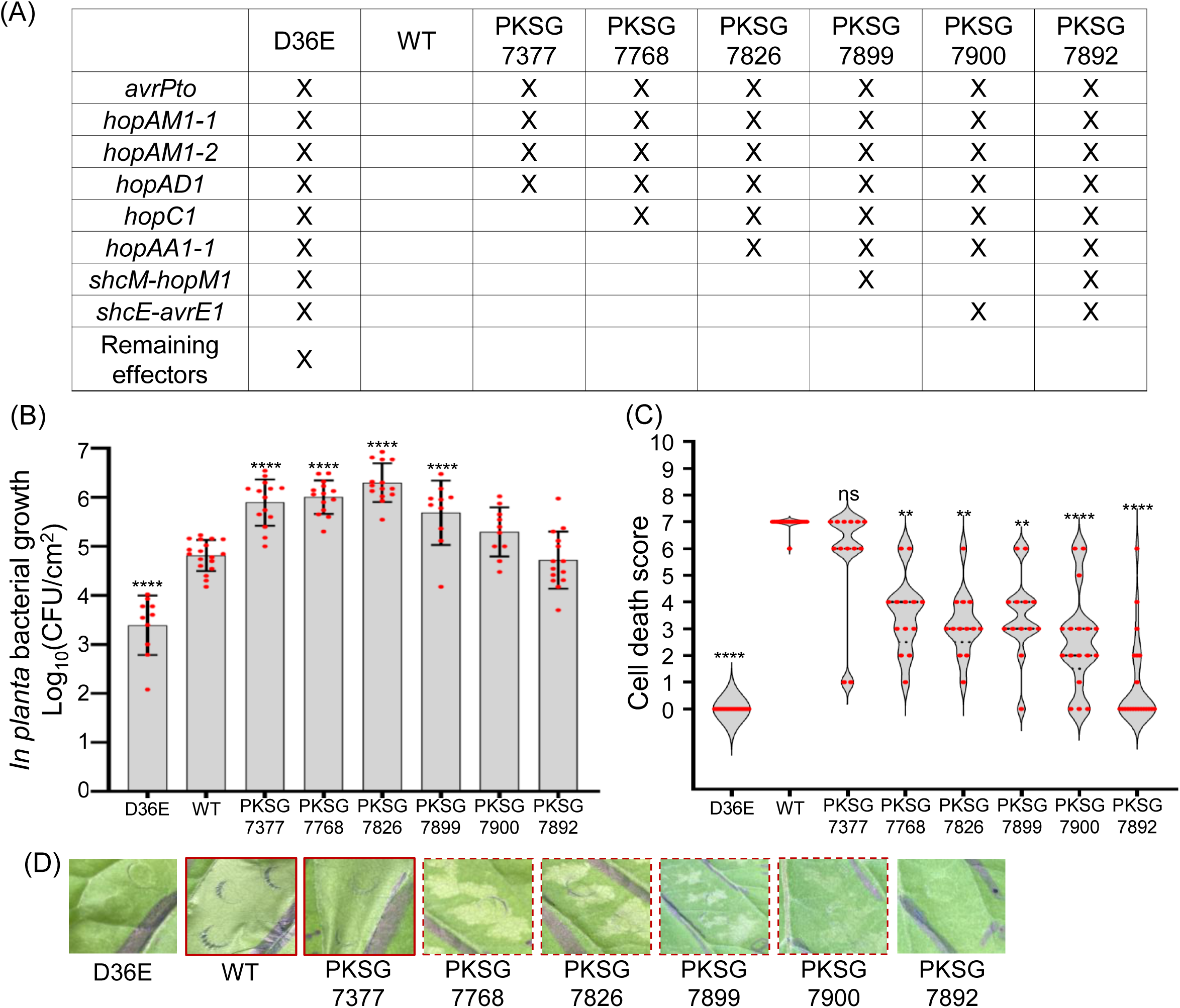
*Pto* DC3000 mutants lacking HR-triggering effectors show enhanced bacterial growth and reduced hypersensitive response phenotypes. **(A)** Table lists mutant names and deleted effectors in each mutant. Effector deletions were performed sequentially and detailed method for effector deletion is explained in Methods section. **(B)** PKSG 7826, lacking *avrPto*, *hopAM1*, *hopAD1*, *hopC1,* and *hopAA1-1* shows enhanced growth compared to *Pto* DC3000 wild-type. Bacterial strains were infiltrated at OD_600nm_=0.0001 using a needleless syringe into SP2273 leaves. *In planta* bacterial growth was quantified four days post infiltration. Individual data points are represented by red dots, with bars indicating the standard deviation. Statistical analysis was conducted using one-way ANOVA followed by Dunnett’s multiple comparison tests (ns: nonsignificant, **: p<0.01, ****: p<0.0001). GRAPHPAD PRISM v.10.0.1 was used for statistical tests. Raw data of bacterial growth are shown in Table S8. **(C)** PKSG 7892, lacking all HR-triggering effectors, does not trigger HR. Bacterial strains were infiltrated at OD_600nm_=0.1 using a needleless syringe into SP2273 leaves. HR was scored one day post infiltration on a scale from 0 (no HR) to 7 (full HR), represented as violin plots. HR scoring criteria are shown in Figure S2, revised from the previous study (Ahn *et al*., 2023). Individual data points are shown as red dots. Statistical significance was conducted using the Kruskal-Wallis test followed by Dunn’s multiple comparison test (ns: nonsignificant, **: p<0.01, ****: p<0.0001), using GRAPHPAD PRISM v.10.01. The raw data of HR scoring are shown in Table S9. **(D)** Representations of HR phenotypes. Bacterial strains were infiltrated at OD_600nm_=0.1 using a needleless syringe into SP2273 leaves. The HR photographs were taken one day post infiltration. The red solid borders indicate the presence of HR, and the red dashed borders indicate weak HR.

To investigate the effect of these additional effector deletions on virulence, we conducted *in planta* bacterial growth assays in SP2273. Infection conditions were identical as described in Figure 3B. Similar to PKSG 7377, PKSG 7768, PKSG 7826, and PKSG 7899 mutants showed significant increase in growth compared to *Pto* DC3000 wild-type (Figure 5B). However, *in planta* growth of PKSG 7900 and PKSG 7892 lacking *shcE-avrE1* was not significantly different from *Pto* DC3000 wild-type (Figure 5B). Next, we tested the HR-inducing activity of these mutant strains in SP2273. Interestingly, PKSG 7768, PKSG 7826, PKSG 7899, and PKSG 7900 triggered significantly reduced HR compared to *Pto* DC3000 wild-type or PKSG 7377 (Figure 5C and 5D). Notably, HR was completely abolished in PKSG 7892, which lacks all effectors previously shown to induce HR in agroinfiltration assay. Finally, we conducted an additional infection assay to monitor disease symptoms caused by effector knockout mutant strains. When SP2273 leaves were syringe-infiltrated with a low concentration of bacterial inoculum (OD_600nm_=0.00001), PKSG 7826 caused visible bacterial speck symptoms (Figure 6A). PKSG 7899 caused a weaker, yet still notable symptoms, while other strains did not produce visible disease symptoms. We further confirmed this phenotype by dip-inoculation assays. We dip-inoculated SP2273 plants with PKSG 7826 (OD_600nm_=0.001). PKSG 7826 showed a mild bacterial speck disease symptom, while *Pto* DC3000 wild-type did not cause visible disease symptoms under the same conditions (Figure 6B). In summary, these results demonstrate that HopC1, HopAA1-1, HopM1, and AvrE1 are required for *Pto* DC3000 avirulence in *S. americanum*. Moreover, despite their HR-inducing activity, HopM1 and AvrE1 appear to carry strong virulence functions as deletion of these effectors significantly reduced *in planta* bacterial growth.

**Figure 6.**
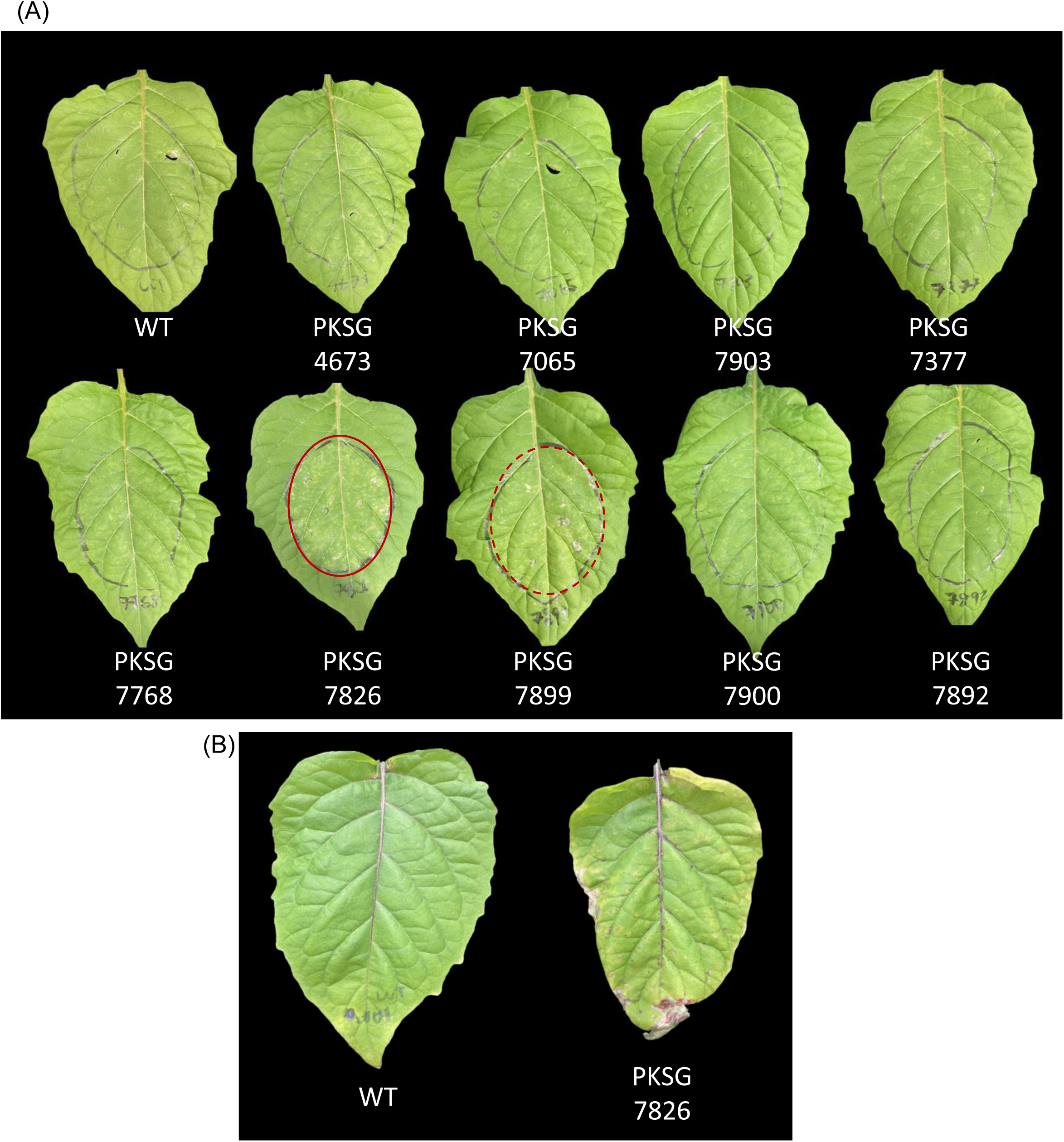
PKSG 7826 causes visible disease symptoms in *Solanum americanum.* **(A)** Bacterial speck phenotype was caused by PKSG 7826. The bacterial inoculum was infiltrated using a needleless syringe into SP2273 leaves. The concentration of bacterial inoculum was OD_600nm_=0.00001. Photographs were taken 7 days post infiltration. *Pto* DC3000 wild-type was used as a negative control. The red border indicates the bacterial disease phenotype and the red dashed border indicates a weak disease phenotype in the infiltrated area. **(B)** PKSG 7826 induces the bacterial speck in SP2273. The leaves were dipped and swirled in the bacterial inoculum for 2 minutes. The OD_600nm_ of bacterial inoculum was 0.001 and mixed with 0.05 % silwet L-77. The photographs were taken 12 days after dipping.

### Conservation of avirulence effectors across multiple *Pseudomonas syringae* strains

Identification of the NLR genes that recognize broadly conserved effectors is considered to provide a source of durable disease resistance. To survey the conservation of seven avirulence effectors identified in this study, we analyzed their presence/absence polymorphism in 117 phytopathogenic *Pseudomonas* strains with available genome sequences in the NCBI database, using effector references from the PsyTEC library (Laflamme et al., 2020). The number of *Pseudomonas* species or pathovar strains analyzed in this research is shown in Table S10. Proteins with an E-value above 1e^-24^ or those with less than 60 % sequence coverage as compared to the effector alleles in PsyTEC database were considered absent (Figure 7A). Based on this criterion, HopAD1, HopAM1, AvrPto, HopC1, HopAA1, HopM1, and AvrE1 were present in 8 (6.8%), 25 (21.4%), 34 (29.1%), 14 (12%), 70 (59.8%), 75 (64.1%) and 109 (93.2%) out of 117 strains, respectively (Figure 7B).

**Figure 7.**
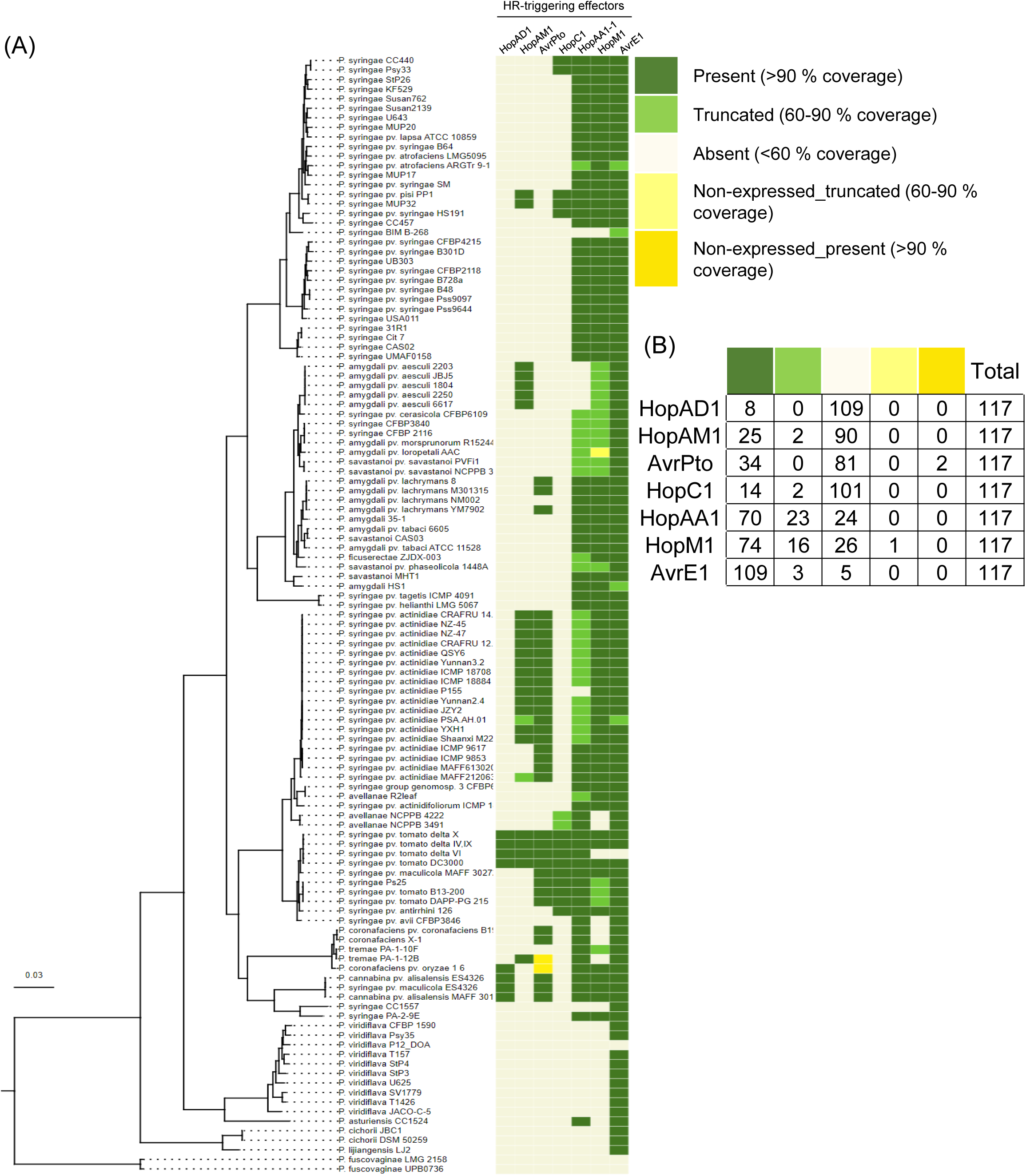
Conservation of hypersensitive response-triggering effectors in *Pseudomonas syringae* strains. **(A)** Each effector sequence was compared to the reference effector sequences from *Pto* DC3000. ‘Non-expressed’ effectors are categorized based on expression information from a previous study (Laflamme et al., 2020). ‘Absent’ indicates effectors with an e-value higher than 1e^-24^ or coverage below 60 %. ‘Truncated’ means each strain’s effector coverage compared to the *Pto* DC3000 is between 60-90 %. ‘Present’ means each strain’s effector and *Pto* DC3000 effector sequence showed over 90 % identity. The number below the table indicates the percentage of strains number having ‘present’ effector genes out of a total of 117 strains. **(B)** Number of *P. syringae* strains number according to their effector presence. The color information is the same as the index in Figure 7A. ‘*P*’ indicates *Pseudomonas* and ‘*Ps*’ indicates *Pseudomonas syringae*.

## DISCUSSION

### Plant immunity triggered by multiple effectors is involved in nonhost resistance

To better understand the interaction between *Solanum americanum* and pathogenic *Pseudomonas syringae* strains, we focused on the well-characterized strain, *Pto* DC3000. Unlike other pathogens such as *P. infestans* and *R. pseudosolanacearum* (Moon *et al*., 2021; Witek *et al*., 2016), *Pto* DC3000 induced defense responses in all tested *S. americanum* accessions (Table 1). Furthermore, well-conserved *P. syringae* effectors such as HopAA1-1, HopM1, and AvrE1 activated defense responses in *S. americanum,* suggesting that *S. americanum* may be a nonhost plant species to *P. syringae*.

The molecular basis of nonhost resistance is still not fully understood, and it is thought to involve multiple contributing factors (Panstruga and Moscou, 2020). One proposed mechanisms is the recognition of pathogen effectors by corresponding NLR genes. For instance, *Phytophthora sojae* is typically a non-adapted pathogen to *N. benthamiana*. However, deletion of AvrNb, an effector recognized by the immune receptor NbPrf, enables *P. sojae* to infect *N. benthamiana* (Dong et al., 2025). This highlights the role of ETI in nonhost resistance. Consistent with our results, previous studies have shown that nonhost resistance can be mediated by the recognition of multiple pathogen effectors. For example, Cevik et al. identified *Albugo candida*-susceptible transgressive segregated lines by using *Arabidopsis thaliana* multiparent advanced generation intercross (MAGIC) lines. Their findings highlighted that nonhost resistance in *A. thaliana* is polygenic and induced through the recognition of multiple *Albugo candida* effectors (Cevik et al., 2019). Similarly, studies in pepper (*Capsicum annuum*) also support this concept. Lee et al. identified several RxLR effectors from *P. infestans* that triggered HR in diverse pepper accessions. It suggests that recognition of multiple effectors contributes to nonhost resistance (Lee et al., 2014). Also, Oh et al., showed that stacked NLR genes in pepper mediate nonhost resistance by recognizing distinct effectors (Oh et al., 2023). While these studies demonstrate nonhost resistance from the plant’s perspective, our research provides additional evidence from the pathogen’s perspective. Specifically, we show that the deletion of multiple HR-triggering effectors enables previously nonpathogenic strain to cause disease in *S. americanum*. Therefore, our research supports the idea that multiple ETIs is one of the genetic bases of nonhost resistance.

### HopM1 and AvrE1 are highly conserved among *Pseudomonas* strains and critical for virulence in *Solanum americanum*

In this study, we show that HopM1 and AvrE1 are crucial for the full virulence of *Pto* DC3000 in *S. americanum*. Previously, multiple studies showed the importance of HopM1 and AvrE1 in pathogen virulence. For instance, deletion of *hopM1* and *avrE1* in *Pto* DC3000 reduces growth and lesion formation in tomato (Badel *et al*., 2006). In *P. syringae* pv. *actinidae*, although HopM1 is non-functional due to the loss of function mutation in *schM*, AvrE1 significantly contributes to virulence in kiwifruit (Jayaraman *et al*., 2020). Furthermore, DspA/E and WtsE, belonging to the AvrE family, are crucial for the full virulence of *Erwinia amylovora* and *Pantoea stewartii*, respectively (Degrave et al., 2015). More recently, AvrE1 and HopM1 were shown to be critical for bacterial virulence in spinach (Mendel et al., 2024). Interestingly, HopM1 and AvrE1 were shown to be critical for the development of water-soaking symptoms during bacterial infection (Xin *et al*., 2016). HopM1 and AvrE1 induce stomatal closure by activating abscisic acid (ABA) signaling, generating an aqueous environment in the apoplast that is favorable for bacterial proliferation (Roussin-Leveillee et al., 2022). Consistent with these findings, our results support the virulence function of HopM1 and AvrE1 in *S. americanum*. It is conceivable that other effectors in *Pto* DC3000 suppress the avirulence activity of HopM1 and AvrE1 in *S. americanum*. It was shown that immune responses mediated by conserved effectors such as HopAA1, HopM1, and AvrE1 can be suppressed by the function of other effectors (Wei et al., 2007). For example, HopI1 suppresses cell death triggered by AvrE1, HopM1, HopQ1-1, HopR1, or HopAM1 in *N. benthamiana* (Wei et al., 2018). Thus, this suggests that these HopM1 and AvrE1 might be indispensable for the full virulence of *Pto* DC3000, although they have the disadvantage of being recognized by unknown plant immune receptors in *S. americanum.* While HopM1 and AvrE1 have been described to have functional redundancy (Kvitko *et al*., 2009), our results show that deletion of either effector leads to reduced bacterial growth in *S. americanum*. This discrepancy may reflect differences in the host resistance gene repertoire or other deleted effectors could influence different levels of redundancy.

### Identifying NLR genes that recognize HR-triggering effectors may enable us to develop durable bacterial speck disease resistant Solanaceae crops

Nonhost resistance confers broad and durable resistance against pathogens (Fonseca and Mysore, 2019). In this study, we hypothesized that nonhost resistance in *S. americanum* is mediated by multiple ETIs. Therefore, identifying NLR genes that recognize effectors involved in nonhost resistance may provide tools for developing durable *P. syringae*-resistance in Solanaceae crops. Several avirulence effectors identified in this study also trigger immune responses in other plant species. For example, HopAD1 induces immune-associated cell death in *N. benthamiana* (Wei *et al*., 2015). AvrPto is known to bind to Pto kinase, activating Prf-dependent disease resistance in tomato. We identified Pto and Prf homologs in the *S. americanum* SP2273 genome, suggesting that recognition of AvrPto in this accession may occur via a similar mechanism with tomato. Furthermore, HopAM1 from *P. syringae* pv. *actinidae* which shares 98.9 % amino acid identity with *Pto* DC3000 allele also induces cell death in *Nicotiana* species, and the homolog from *P. syringae* pv. *pisi* induces cell death in pea cultivars (Choi et al., 2017; Cournoyer et al., 1995; Eastman et al., 2022). HopAM1 contains a Toll/interleukin-1 receptor (TIR) domain such as TIR-type NLR, which activates immune signaling and cell death in plants (Eastman *et al*., 2022). Therefore, HopAM1 may trigger cell death independently of a specific resistance gene. HopC1 homolog from *P. syringae* pv. *pisi* which shares 99.6 % amino acid identity with *Pto* DC3000 allele, acts as an avirulence effector in bean (Arnold et al., 2001; Baltrus et al., 2012). HopM1 and AvrE1 also trigger cell death in *N. benthamiana* (Wei et al., 2018). Despite various research on HR phenotypes and immune responses for theses effectors, resistance genes recognizing these avirulence effectors are not well-studied in Solanaceae plants. In *Arabidopsis*, the immune receptor, CEL-ACTIVATED RESISTANCE1 (CAR1), recognizes AvrE1 and HopAA1-1 (Laflamme *et al*., 2020). However, an NCBI BLAST search using the CAR1 sequence (AT1G50180.1) found no clear homolog in the SP2273 genome (data not shown). This could be an example of convergent evolution, where an effector protein is recognized by resistance genes with no sequence similarity in different plant species (Kim et al., 2023). In the future, NLR genes recognizing avirulence *Pto* DC3000 effectors could be identified through natural variations of *S. americanum* accessions. In particular, NLRs that detect conserved avirulence effectors such as AvrE1, HopM1, and HopAA1 may be especially valuable for developing durable resistance against *P. syringae* in Solanaceae crops.

### Virulent *Pto* DC3000 mutant strain is a valuable tool for studying effector-triggered immunity in *Solanum americanum*

*S. americanum* is a promising model Solanaceae plant for identifying resistance genes against diverse phytopathogens (Witek *et al*., 2016). However, the lack of virulent strains for *S. americanum* makes it challenging to study the interaction between effectors and host plants. In this study, we generated virulent strains using the model bacterial pathogen *Pto* DC3000. In *N. benthamiana*, a well-established model Solanaceae plant, *Pto* DC3000 Δ*hopQ1-1* is commonly used for *in planta* bacterial growth assays (Wei *et al*., 2007). Similarly, we can use the virulent PKSG 7826 strain to perform *in planta* bacterial growth assays in *S. americanum*. Thus, this study highlights the potential of *S. americanum* to serve as a model Solanaceae plant for ETI studies alongside *N. benthamiana*.

## METHODS

### Plant growth conditions

*Solanum americanum* and *Nicotiana benthamiana* plants were grown in Baroker soil mix (4 % zeolite, 7 % perlite, 6 % vermiculite, 68 % cocopeat, 14.73 % peat moss; Seoulbio, http://www.seoulbio.co.kr) at 23 ℃ with 11 hours of light per day. *S. americanum* was used for agroinfiltration and *Pseudomonas syringae* pv. *tomato* DC3000 *in planta* growth assays. *N. benthamiana* was used for agroinfiltration followed by total protein extraction and immunoblot analysis.

### Bacterial strains and culture conditions

The bacterial strains used in this study are listed in Table 2. *Escherichia coli* DH5α and *Agrobacterium tumefaciens* AGL1 strains were cultured in Luria-Bertani (LB) broth containing appropriate antibiotics. *Pseudomonas syringae* strains were grown on King’s B (KB) medium with appropriate antibiotics. Antibiotics for bacterial strains are shown in Table S2. Antibiotic concentrations used were: Carbenicillin (100 μg/mL), Kanamycin (50 μg/mL), Gentamycin (20 μg/mL), Rifampicin (50 μg/mL), Spectinomycin (100 μg/mL). *E. coli* was grown at 37 ℃, and *A. tumefaciens* and *P. syringae* were grown at 28 ℃.

### Construction of plasmids

To clone *Pto* DC3000 type III effector genes, we used *Pto* DC3000 chromosome data (GenBank: AE016853.1) and pDC3000a plasmid data (NCBI: NC_004633.1). If effector sequences are over 2000 bp, we divided them into modules (about 500 - 1500 bp) for efficient golden-gate (GG) assembly. First, *BsaI* site flanked nucleotide sequences of *hopA1*, *hopAD1*, *hopAF1* (*BsaI* mutagenized), *hopAM1*, *hopB1* (*BsaI* mutagenized), *hopE1*, *hopI1* (codon optimized), *hopK1*, *hopY1*, *hopD1*_module2 (*BsaI* mutagenized), *hopR1*_module6 (*BsaI* mutagenized), *hopN1* (*BsaI* mutagenized), *hopAA1-2*_module2 (*BsaI* mutagenized), *avrE1*_module 4 (*BsaI* mutagenized) were synthesized (Twist Biosciences, South San Francisco, CA, USA). Other effectors or effector modules were PCR amplified. *Pto* DC3000 genomic DNA (extracted using Wizard Genomic DNA purification kit; Promega, https://www.promega.com/) was used as the PCR template. All DNA fragments were flanked by *BsaI* restriction enzyme site and GG overhangs to make GG modules. Effectors were cloned into pICH41021 vector (*BsaI* mutagenized pUC19, hereafter pUC19B) by blunt-end ligation using *SmaI* or *Eco53KI* restriction enzyme. The pUC19B modules were cloned into the binary vector (pICH86988) with a C-terminal 6x HA epitope tag using golden gate assembly using *BsaI* restriction enzyme (Engler et al., 2008).

For generating *Pto* DC3000 type III effector knockout plasmids, the upstream and downstream of the target effector region (around 1∼1.8 kb) were PCR-amplified. DNA fragments were flanked by *BsaI* restriction sites and GG overhangs. The amplified PCR templates were cloned into pUC19B vector. The upstream and downstream pUC19B modules were assembled into suicide pK18mobsacB-GG vector via GG assembly using *BsaI* restriction enzyme (Jayaraman *et al*., 2020). pK18mobsacB-GG vector was derived by removing the multiple cloning site (MCS) of the original pk18mobsacB vector. The removed MCS was replaced with a *BsaI* restriction site flanked MCS of pICH86988 (Jayaraman *et al*., 2020). The primers or synthesized sequences for pUC19B module cloning are listed in Table S11 and Table S12. All pUC19B modules and GG-assembled constructs were transformed into *E. coli* DH5α using electroporation method (1.8 kV pulse using 1 mm electroporation cuvette). *E. coli* transformants were selected on LB media containing appropriate antibiotics for destination vectors (see Table S3 for details). The insert sequences of pUC19B modules were validated using Sanger sequencing and GG-assembled constructs were verified using restriction enzyme digestion.

### *Agrobacterium*-mediated transient transformation

All effectors used in this study were cloned in the binary vector pICH86988 for *Agrobacterium*-mediated transient transformation (agroinfiltration). These binary constructs were transformed into *A. tumefaciens* AGL1 strain using electroporation (2.2 kV pulse using 1mm electroporation cuvette). AGL1 transformants were selected on LB media containing Carbenicillin (100 μg/mL) and Kanamycin (50 μg/mL). For agroinfiltration, AGL1 strains carrying effector constructs and P19, a viral suppressor of RNA silencing, were grown in liquid LB with appropriate antibiotics. The cultured cells were resuspended in *Agrobacterium* infiltration buffer (10 mM MgCl_2_ and 10 mM MES (pH 5.6)) and diluted to OD_600nm_ 0.2 (P19) to 0.4 (effector). The diluted *Agrobacterium* inoculums carrying effector constructs and P19 were mixed and infiltrated using a 1 mL needleless syringe into plant leaves.

### Immunoblot analysis

The total protein was extracted from six leaf discs (8mm diameter) taken from the infiltrated area of *N. benthamiana* two days post infiltration. The leaf discs were snap-frozen in liquid nitrogen and ground with 200 μL of 5X SDS protein loading buffer (250mM Tris-HCl (pH 6.8), 8 % sodium dodecyl sulfate (SDS), 0.1 % Bromophenol blue, 40 % (v/v) Glycerol, and 100 mM Dithiothreitol). The total protein samples were boiled for 10 minutes at 95 ℃. For SDS page, 20 μL of protein samples were loaded in polyacrylamide gel and ran for 1 hour (130 V). HopA1, HopAD1, HopAF1, HopAM1, HopB1, HopE1, AvrPto, AvrPtoB, HopI1, HopK1, HopY1, HopH1, HopM1, HopN1, HopO1-1, HopQ1-1, HopF2, HopT1-1, HopU1, HopV1, and HopX1 protein samples were loaded in 10 % polyacrylamide gel. HopAA1-1, HopAA1-2, HopAO1, AvrE1, HopC1, HopD1, HopR1, and HopG1 were loaded in 4∼12 % Mini-PROTEAN TGX Precast Gels (Bio-Rad, https://www.bio-rad.com/). Proteins were transferred from the polyacrylamide gel into PVDF membranes for one hour (100 V). Protein-transferred PVDF membranes were blocked using 5 % (w/v) skim milk in Tris-Buffered Saline (pH 7.4), and 0.1 % Tween20 (TBST) for 30 minutes. After blocking, anti-HA (Roche 11867423001; 1:2000 dilution) antibodies were added to the blocking buffer and incubated for 1 hour at room temperature. The membrane was washed for five minutes (five times) using TBST. The membranes were blocked in a blocking buffer for 30 minutes at room temperature. Anti-rat secondary antibody (Sigma A9037; 1:20000 dilution) was added to the blocking buffer. The membranes were incubated for 1 hour 30 minutes at room temperature. Membranes were washed for five minutes (five times) using TBST. Proteins were visualized using Super Signal West Pico and Femto Chemiluminescent substrate (Thermo Fisher Scientific, https://www.thermofisher.com/) through ChemiDoc XRS+ with Image Lab Software (Bio-Rad). After visualization, Membranes were stained with Ponceau S to estimate the quantity of proteins.

### Hypersensitive response, *in planta* growth, and dip inoculation assays using *Pseudomonas*

*Pseudomonas syringae* pv. *tomato* DC3000 wild-type and mutant strains were grown on KB agar with appropriate antibiotics. *Pto* DC3000 cells were resuspended in autoclaved *Pseudomonas* infiltration buffer (10 mM MgCl_2_). The inoculum was diluted to OD_600nm_=0.1 for HR assays, OD_600nm_=0.0001 for *in planta* growth assays and OD_600nm_=0.001 for dip inoculation. The bacterial inoculum was infiltrated using a needleless syringe into four to five-week-old *S. americanum* leaves. HR was scored from 0 to 7 one day post infiltration. HR scoring criteria were described in Figure S2 and revised from the previous study (Ahn *et al*., 2023). The HR scores triggered by *Pto* DC3000 wild-type and effector knockout mutants are shown in the violin plots showing individual replicate data. For *in planta* bacterial growth, two leaf discs (8 mm diameter) were grounded and diluted in 500 μL of autoclaved *Pseudomonas* infiltration buffer. Serial dilutions (10 μL) were spotted on KB agar with appropriate antibiotics. Colonies were counted after two days at 28 ℃. Three to four plants were used per batch repeat. The bacterial growth (CFU/cm^2^) is shown in the bar graph with individual data. For dip inoculation, the bacteria inoculum was diluted to OD_600nm_=0.001 in *Pseudomonas* infiltration buffer and 0.05 % Silwet L-77. Four- to five-week-old SP2273 leaves were dipped and gently swirled for 2 minutes. The dipped *S. americanum* leaves were covered for one day to maintain humidity (11h light, 23 ℃). The photographs were taken 12 days after dip inoculation.

### Generation of *Pseudomonas syringae* pv. *tomato* DC3000 effector knock-out strains

Details of effector deletion in *Pto* DC3000 can be found in (Jayaraman *et al*., 2020). Recipient bacteria were incubated on KB agar with Rifampicin (50 μg/mL). Helper *E. coli* HB101 and Donor *E. coli* DH5α carrying suicide vector (pK18mobsacB-GG) containing upstream and downstream regions of the target effector were cultured on LB medium containing Kanamycin (50 μg/mL). All three bacteria strains were mixed on LB agar without antibiotics and incubated for 6-7 hours at 28 ℃. Mixed bacteria were streaked on KB agar with Rifampicin (50 μg/mL) and Kanamycin (50 μg/mL). After 2 days of incubation at 28 ℃, a single colony was selected. A single colony was inoculated into liquid LB media with Rifampicin (50 μg/mL) and grown for one day at 28 ℃. The cultured bacteria were streaked on KB agar media containing Rifampicin (50 μg/mL) and 10 % sucrose (w/v). After two days of incubation at 28 ℃, a single colony was selected. Successful knockouts showed reduced band sizes using specific PCR primers (see Table S11 for details).

## DATA AVAILABILITY

All relevant data can be found in the manuscript, Supplemental information, and public databases (NCBI).

## AUTHOR CONTRIBUTIONS

JK and KHS designed the experiments. JK performed the experiments. JK, MVZ, and KHS analyzed the data. JK and KHS wrote the manuscript.

## ACKNOWLEDGEMENTS

We thank Prof. Duck Hwan Park (Kangwon National University, Republic of Korea) and Dr. Jay Jayaraman (Plant and Food Research, New Zealand) for sharing materials. The authors declare no competing interests.

## SUPPLEMENTAL INFORMATION

**Figures S1.** Generation and confirmation of primary candidate effectors knockout, related to Figure 3

**Figure S2.** Hypersensitive response scoring criteria in *Solanum americanum,* Related to Figure 3 and Figure 5

**Figure S3.** Generation and confirmation of effector knockout, Related to Figure 5

**Table S1.** *Solanum americanum* accessions used in this study

**Table S2.** Bacterial strains used in this study

**Table S3.** Plasmids used in this study

**Table S4.** Raw data of *in planta* bacterial growth, Related to Figure 1

**Table S5.** Type III effector expected protein size, Related to Figure 2 and Figure 4

**Table S6.** Raw data of *in planta* bacterial growth, Related to Figure 3

**Table S7.** Raw data of hypersensitive response scoring, Related to Figure 3

**Table S8.** Raw data of *in planta* bacterial growth, Related to Figure 5

**Table S9.** Raw data of hypersensitive response scoring, Related to Figure 5

**Table S10.** Numbers of species or pathovars analyzed in this study, Related to Figure 7

**Table S11.** Excel file containing information of primers used in this study

**Table S12.** Excel file containing sequence information of synthesized constructs

## REFERENCES

Ahn, Y.J., Kim, H., Choi, S., Mazo-Molina, C., Prokchorchik, M., Zhang, N., Kim, B., Mang, H., Koehler, N., and Kim, J. (2023). Ptr1 and ZAR1 immune receptors confer overlapping and distinct bacterial pathogen effector specificities. New Phytologist 239:1935–1953.

Alfano, J.R., Charkowski, A.O., Deng, W.L., Badel, J.L., Petnicki-Ocwieja, T., van Dijk, K., and Collmer, A. (2000). The Pseudomonas syringae Hrp pathogenicity island has a tripartite mosaic structure composed of a cluster of type III secretion genes bounded by exchangeable effector and conserved effector loci that contribute to parasitic fitness and pathogenicity in plants. Proc Natl Acad Sci U S A 97:4856–4861. 10.1073/pnas.97.9.4856.

Arnold, D.L., Jackson, R.W., Fillingham, A.J., Goss, S.C., Taylor, J.D., Mansfield, J.W., and Vivian, A. (2001). Highly conserved sequences flank avirulence genes: isolation of novel avirulence genes from Pseudomonas syringae pv. pisi. Microbiology 147:1171–1182.

Arora, S., Steuernagel, B., Gaurav, K., Chandramohan, S., Long, Y., Matny, O., Johnson, R., Enk, J., Periyannan, S., Singh, N., et al. (2019). Resistance gene cloning from a wild crop relative by sequence capture and association genetics. Nat Biotechnol 37:139–143. 10.1038/s41587-018-0007-9.

Badel, J.L., Shimizu, R., Oh, H.-S., and Collmer, A. (2006). A Pseudomonas syringae pv. tomato avrE1/hopM1 mutant is severely reduced in growth and lesion formation in tomato. Molecular plant-microbe interactions 19:99–111.

Badel, J.L., Nomura, K., Bandyopadhyay, S., Shimizu, R., Collmer, A., and He, S.Y. (2003). Pseudomonas syringae pv. tomato DC3000 HopPtoM (CEL ORF3) is important for lesion formation but not growth in tomato and is secreted and translocated by the Hrp type III secretion system in a chaperone-dependent manner. Molecular microbiology 49:1239–1251.

Baltrus, D.A., Nishimura, M.T., Dougherty, K.M., Biswas, S., Mukhtar, M.S., Vicente, J., Holub, E.B., and Dangl, J.L. (2012). The molecular basis of host specialization in bean pathovars of Pseudomonas syringae. Molecular plant-microbe interactions 25:877–888.

Bi, G., Su, M., Li, N., Liang, Y., Dang, S., Xu, J., Hu, M., Wang, J., Zou, M., Deng, Y., et al. (2021). The ZAR1 resistosome is a calcium-permeable channel triggering plant immune signaling. Cell 184:3528–3541 e3512. 10.1016/j.cell.2021.05.003.

Buell, C.R., Joardar, V., Lindeberg, M., Selengut, J., Paulsen, I.T., Gwinn, M.L., Dodson, R.J., Deboy, R.T., Durkin, A.S., Kolonay, J.F., et al. (2003). The complete genome sequence of the Arabidopsis and tomato pathogen Pseudomonas syringae pv. tomato DC3000. Proc Natl Acad Sci U S A 100:10181–10186. 10.1073/pnas.1731982100.

Cevik, V., Boutrot, F., Apel, W., Robert-Seilaniantz, A., Furzer, O.J., Redkar, A., Castel, B., Kover, P.X., Prince, D.C., and Holub, E.B. (2019). Transgressive segregation reveals mechanisms of Arabidopsis immunity to Brassica-infecting races of white rust (Albugo candida). Proceedings of the National Academy of Sciences 116:2767–2773.

Choi, S., Jayaraman, J., Segonzac, C., Park, H.-J., Park, H., Han, S.-W., and Sohn, K.H. (2017). Pseudomonas syringae pv. actinidiae type III effectors localized at multiple cellular compartments activate or suppress innate immune responses in Nicotiana benthamiana. Frontiers in Plant Science 8:309600.

Cournoyer, B., Sharp, J., Astuto, A., Gibbon, M., Taylor, J., and Vivian, A. (1995). Molecular characterization of the Pseudomonas syringae pv. pisi plasmid-borne avirulence gene avrPpiB which matches the R3 resistance locus in pea. Molecular Plant-microbe Interactions: MPMI 8:700–708.

Cunnac, S., Chakravarthy, S., Kvitko, B.H., Russell, A.B., Martin, G.B., and Collmer, A. (2011). Genetic disassembly and combinatorial reassembly identify a minimal functional repertoire of type III effectors in Pseudomonas syringae. Proceedings of the National Academy of Sciences 108:2975–2980.

Cuppels, D.A. (1986). Generation and characterization of Tn 5 insertion mutations in Pseudomonas syringae pv. tomato. Applied and environmental microbiology 51:323–327.

Dangl, J.L., Horvath, D.M., and Staskawicz, B.J. (2013). Pivoting the Plant Immune System from Dissection to Deployment. Science 341:746–751. doi:10.1126/science.1236011.

Degrave, A., Siamer, S., Boureau, T., and Barny, M.A. (2015). The AvrE superfamily: ancestral type III effectors involved in suppression of pathogen-associated molecular pattern-triggered immunity. Mol Plant Pathol 16:899–905. 10.1111/mpp.12237.

Dong, X., Lu, X., Zhu, H., Zhu, Z., Ji, P., Li, X., Li, T., Zhang, X., Ai, G., and Dou, D. (2025). A typical NLR recognizes a family of structurally conserved effectors to confer plant resistance against adapted and non-adapted Phytophthora pathogens. Molecular Plant.

Eastman, S., Smith, T., Zaydman, M.A., Kim, P., Martinez, S., Damaraju, N., DiAntonio, A., Milbrandt, J., Clemente, T.E., Alfano, J.R., and Guo, M. (2022). A phytobacterial TIR domain effector manipulates NAD+ to promote virulence. New Phytologist 233:890–904. 10.1111/nph.17805.

Engler, C., Kandzia, R., and Marillonnet, S. (2008). A one pot, one step, precision cloning method with high throughput capability. PloS one 3:e3647.

Figurski, D.H., and Helinski, D.R. (1979). Replication of an origin-containing derivative of plasmid RK2 dependent on a plasmid function provided in trans. Proc Natl Acad Sci U S A 76:1648–1652. 10.1073/pnas.76.4.1648.

Fonseca, J.P., and Mysore, K.S. (2019). Genes involved in nonhost disease resistance as a key to engineer durable resistance in crops. Plant Sci 279:108–116. 10.1016/j.plantsci.2018.07.002.

Gill, U.S., Lee, S., and Mysore, K.S. (2015). Host versus nonhost resistance: distinct wars with similar arsenals. Phytopathology 105:580–587. 10.1094/PHYTO-11-14-0298-RVW.

Jayaraman, J., Yoon, M., Applegate, E.R., Stroud, E.A., and Templeton, M.D. (2020). AvrE1 and HopR1 from Pseudomonas syringae pv. actinidiae are additively required for full virulence on kiwifruit. Molecular Plant Pathology 21:1467–1480. 10.1111/mpp.12989.

Jayaraman, J., Choi, S., Prokchorchik, M., Choi, D.S., Spiandore, A., Rikkerink, E.H., Templeton, M.D., Segonzac, C., and Sohn, K.H. (2017). A bacterial acetyltransferase triggers immunity in Arabidopsis thaliana independent of hypersensitive response. Scientific Reports 7:3557.

Jones, J.D., and Dangl, J.L. (2006). The plant immune system. Nature 444:323–329. 10.1038/nature05286.

Kim, B., Choi, J., and Segonzac, C. (2022). Tackling multiple bacterial diseases of Solanaceae with a handful of immune receptors. Horticulture, Environment, and Biotechnology 63:149–160. 10.1007/s13580-021-00415-1.

Kim, H., Ahn, Y.J., Lee, H., Chung, E.H., Segonzac, C., and Sohn, K.H. (2023). Diversified host target families mediate convergently evolved effector recognition across plant species. Curr Opin Plant Biol 74:102398. 10.1016/j.pbi.2023.102398.

Kourelis, J., and van der Hoorn, R.A.L. (2018). Defended to the Nines: 25 Years of Resistance Gene Cloning Identifies Nine Mechanisms for R Protein Function. Plant Cell 30:285–299. 10.1105/tpc.17.00579.

Kvitko, B.H., Park, D.H., Velásquez, A.C., Wei, C.-F., Russell, A.B., Martin, G.B., Schneider, D.J., and Collmer, A. (2009). Deletions in the repertoire of Pseudomonas syringae pv. tomato DC3000 type III secretion effector genes reveal functional overlap among effectors. PLoS pathogens 5:e1000388.

Laflamme, B., Dillon, M.M., Martel, A., Almeida, R.N., Desveaux, D., and Guttman, D.S. (2020). The pan-genome effector-triggered immunity landscape of a host-pathogen interaction. Science 367:763–768.

Lee, H.A., Kim, S.Y., Oh, S.K., Yeom, S.I., Kim, S.B., Kim, M.S., Kamoun, S., and Choi, D. (2014). Multiple recognition of RXLR effectors is associated with nonhost resistance of pepper against Phytophthora infestans. New Phytol 203:926–938. 10.1111/nph.12861.

Lin, X., Olave-Achury, A., Heal, R., Pais, M., Witek, K., Ahn, H.K., Zhao, H., Bhanvadia, S., Karki, H.S., Song, T., et al. (2022). A potato late blight resistance gene protects against multiple Phytophthora species by recognizing a broadly conserved RXLR-WY effector. Mol Plant 15:1457–1469. 10.1016/j.molp.2022.07.012.

Lin, X., Jia, Y., Heal, R., Prokchorchik, M., Sindalovskaya, M., Olave-Achury, A., Makechemu, M., Fairhead, S., Noureen, A., Heo, J., et al. (2023). Solanum americanum genome-assisted discovery of immune receptors that detect potato late blight pathogen effectors. Nat Genet 55:1579–1588. 10.1038/s41588-023-01486-9.

Lindeberg, M., Cartinhour, S., Myers, C.R., Schechter, L.M., Schneider, D.J., and Collmer, A. (2006). Closing the circle on the discovery of genes encoding Hrp regulon members and type III secretion system effectors in the genomes of three model Pseudomonas syringae strains. Molecular Plant-Microbe Interactions 19:1151–1158.

Macho, A.P. (2016). Subversion of plant cellular functions by bacterial type-III effectors: beyond suppression of immunity. New Phytol 210:51–57. 10.1111/nph.13605.

Mendel, M., Zuijdgeest, X.C.L., Berg, F.v.d., Meer, L.v.d., Elberse, J., Skiadas, P., Seidl, M.F., Ackerveken, G.V.d., and Jonge, R.d. (2024). Exploiting <em>Pseudomonas syringae</em> Type 3 secretion to study effector contribution to disease in spinach. bioRxiv:2024.2006.2014.599008. 10.1101/2024.06.14.599008.

Moon, H., Pandey, A., Yoon, H., Choi, S., Jeon, H., Prokchorchik, M., Jung, G., Witek, K., Valls, M., and McCann, H.C. (2021). Identification of RipAZ1 as an avirulence determinant of Ralstonia solanacearum in Solanum americanum. Molecular plant pathology 22:317–333.

Munkvold, K.R., Russell, A.B., Kvitko, B.H., and Collmer, A. (2009). Pseudomonas syringae pv. tomato DC3000 type III effector HopAA1-1 functions redundantly with chlorosis-promoting factor PSPTO4723 to produce bacterial speck lesions in host tomato. Mol Plant Microbe Interact 22:1341–1355. 10.1094/mpmi-22-11-1341.

Oh, S., Kim, S., Park, H.J., Kim, M.S., Seo, M.K., Wu, C.H., Lee, H.A., Kim, H.S., Kamoun, S., and Choi, D. (2023). Nucleotide-binding leucine-rich repeat network underlies nonhost resistance of pepper against the Irish potato famine pathogen Phytophthora infestans. Plant biotechnology journal 21:1361–1372.

Panstruga, R., and Moscou, M.J. (2020). What is the Molecular Basis of Nonhost Resistance? Mol Plant Microbe Interact 33:1253–1264. 10.1094/MPMI-06-20-0161-CR.

Roussin-Leveillee, C., Lajeunesse, G., St-Amand, M., Veerapen, V.P., Silva-Martins, G., Nomura, K., Brassard, S., Bolaji, A., He, S.Y., and Moffett, P. (2022). Evolutionarily conserved bacterial effectors hijack abscisic acid signaling to induce an aqueous environment in the apoplast. Cell Host Microbe 30:489–501 e484. 10.1016/j.chom.2022.02.006.

Schäfer, A., Tauch, A., Jäger, W., Kalinowski, J., Thierbach, G., and Pühler, A. (1994). Small mobilizable multi-purpose cloning vectors derived from the Escherichia coli plasmids pK18 and pK19: selection of defined deletions in the chromosome of Corynebacterium glutamicum. Gene 145:69–73. 10.1016/0378-1119(94)90324-7.

Schultink, A., Qi, T., Lee, A., Steinbrenner, A.D., and Staskawicz, B. (2017). Roq1 mediates recognition of the Xanthomonas and Pseudomonas effector proteins XopQ and HopQ1. The Plant Journal 92:787–795.

Wang, J., Hu, M., Wang, J., Qi, J., Han, Z., Wang, G., Qi, Y., Wang, H.W., Zhou, J.M., and Chai, J. (2019). Reconstitution and structure of a plant NLR resistosome conferring immunity. Science 36410.1126/science.aav5870.

Weber, E., Engler, C., Gruetzner, R., Werner, S., and Marillonnet, S. (2011). A modular cloning system for standardized assembly of multigene constructs. PloS one 6:e16765.

Wei, C.F., Kvitko, B.H., Shimizu, R., Crabill, E., Alfano, J.R., Lin, N.C., Martin, G.B., Huang, H.C., and Collmer, A. (2007). A Pseudomonas syringae pv. tomato DC3000 mutant lacking the type III effector HopQ1-1 is able to cause disease in the model plant Nicotiana benthamiana. The Plant Journal 51:32–46.

Wei, H.-L., Zhang, W., and Collmer, A. (2018). Modular study of the type III effector repertoire in Pseudomonas syringae pv. tomato DC3000 reveals a matrix of effector interplay in pathogenesis. Cell reports 23:1630–1638.

Wei, H.-L., Chakravarthy, S., Mathieu, J., Helmann, T.C., Stodghill, P., Swingle, B., Martin, G.B., and Collmer, A. (2015). Pseudomonas syringae pv. tomato DC3000 type III secretion effector polymutants reveal an interplay between HopAD1 and AvrPtoB. Cell host & microbe 17:752–762.

Witek, K., Jupe, F., Witek, A.I., Baker, D., Clark, M.D., and Jones, J.D. (2016). Accelerated cloning of a potato late blight-resistance gene using RenSeq and SMRT sequencing. Nat Biotechnol 34:656–660. 10.1038/nbt.3540.

Witek, K., Lin, X., Karki, H., Jupe, F., Witek, A., and Steuernagel, B. (2021). A complex resistance locus in Solanum americanum recognizes a conserved Phytophthora effector. Nat Plants. 2021; 7: 198–208.

Xiang, T., Zong, N., Zou, Y., Wu, Y., Zhang, J., Xing, W., Li, Y., Tang, X., Zhu, L., Chai, J., and Zhou, J.M. (2008). Pseudomonas syringae effector AvrPto blocks innate immunity by targeting receptor kinases. Curr Biol 18:74–80. 10.1016/j.cub.2007.12.020.

Xin, X.-F., and He, S.Y. (2013). Pseudomonas syringae pv. tomato DC3000: a model pathogen for probing disease susceptibility and hormone signaling in plants. Annual review of phytopathology 51:473–498.

Xin, X.F., Kvitko, B., and He, S.Y. (2018). Pseudomonas syringae: what it takes to be a pathogen. Nat Rev Microbiol 16:316–328. 10.1038/nrmicro.2018.17.

Zipfel, C., and Rathjen, J.P. (2008). Plant immunity: AvrPto targets the frontline. Curr Biol 18:R218–220. 10.1016/j.cub.2008.01.016.

