## Supplementary material for "Multiple effectors trigger nonhost resistance in *Solanum americanum* against *Pseudomonas syringae*": Fig S1-S3, Table S1-S10

#### Figure S1, Generation and confirmation of primary candidate effectors knockout, Related to Figure 3

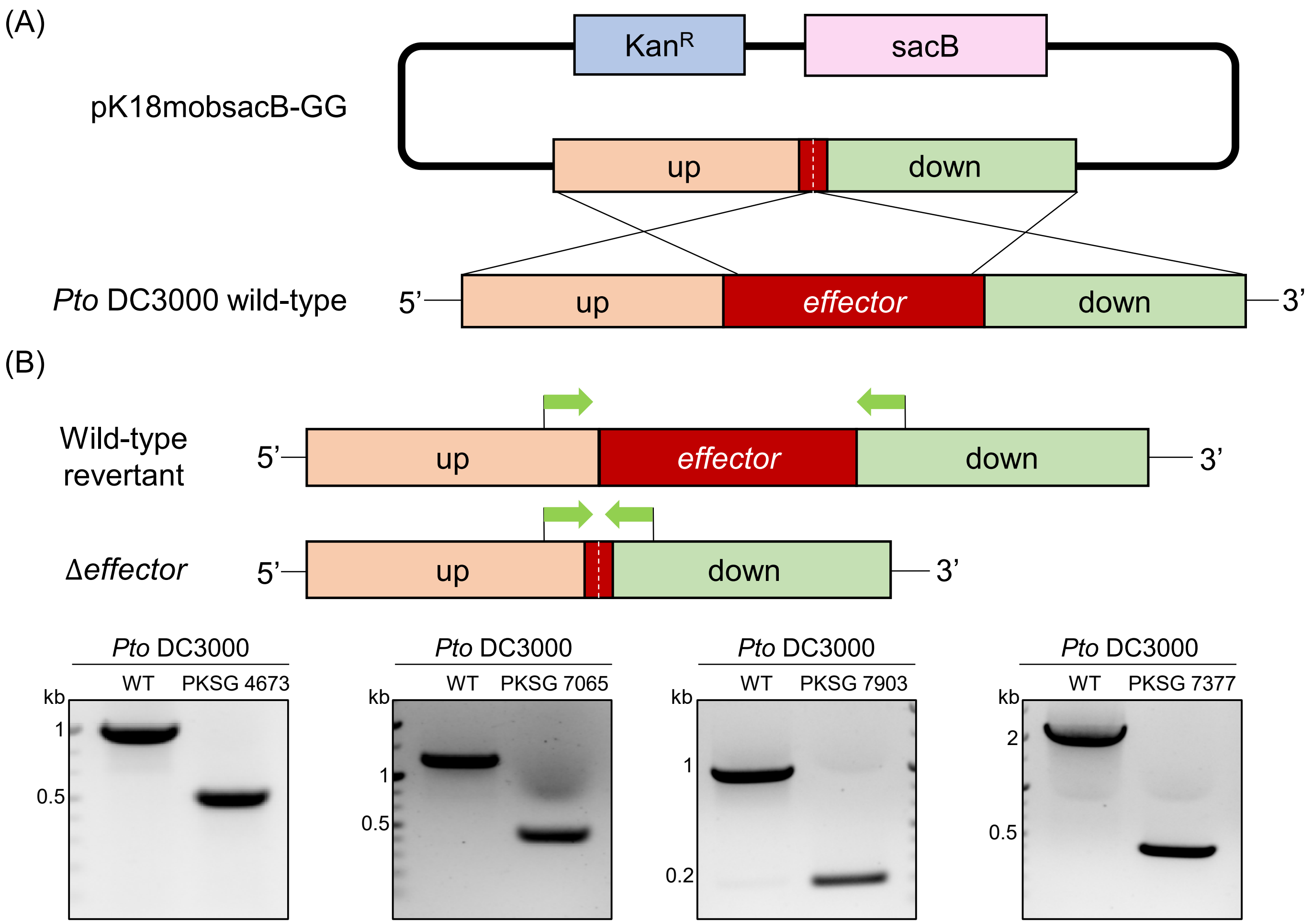


(C)

| | Primer (PKSP) | Expected band size (bp) | |
|---|---|---|---|
| | | Wild-type | Δ*effector* |
| *avrPto* | 3910+6358 | 897 | 148 |
| *hopAM1-1* | 6361+6362 | 1260 | 442 |
| *hopAM1-2* | 5719+5720 | 1008 | 190 |
| *hopAD1* | 6359+6360 | 2295 | 412 |

(A) Schematic representation of effector knockout in *Pto* DC3000. For *avrPto* knockout, the upstream flanking region was designed with 1077 bp and the downstream region with 1521 bp. Both upstream and downstream regions included 6 bp of *avrPto* gene 5' and 3' end, respectively. For *hopAM1-1* knockout, the upstream flanking region of *hopAM1-1* flanking region was 1476 bp, and downstream was 1515 bp. The upstream and downstream regions contained 3 bp and 6 bp of the *hopAM1-1* 5' and 3' end respectively. Similarly, for *hopAM1-2* knockout, the upstream of *hopAM1-2* flanking region was 1497 bp, and downstream was 1500 bp including 3 bp and 6 bp of 5' and 3' end of *hopAM1-2* gene, respectively. For *hopAD1* knockout, the upstream flanking region was 1515 bp and the downstream was 1422 bp with both regions containing 6 bp of the *hopAD1* 5' and 3' end each. The upstream and downstream flanking region constructs are Goldengate cloned into pK18mobsacB-GG vector [S1]. A detailed effector gene knockout method is shown in the method section. (B) PCR confirmation effector knockout in *Pto* DC3000. Primers are designed to be located in the flanking region of the effector gene. The reduced band size indicates the deletion of the gene. Green arrows indicate the primers used to confirm the knockout. (C) The table shows the effector knockouts' primer names and expected band sizes. PKSP refers to the used primer name. The primer sequences are shown in Table S11.

Figure S2. Hypersensitive response scoring criteria in *Solanum americanum*, Related to Figure 3 and Figure 5

| HR score | HR photo | Description |
|---|---|---|
| 0 | 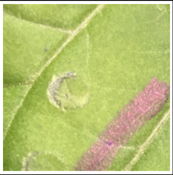 | No cell death is observed |
| 1 | 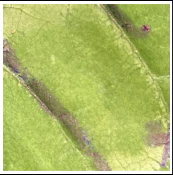 | A small area of cell death is observed at the periphery of the infiltrated area |
| 2 | 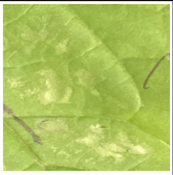 | Small spots of cell death are observed within the infiltration area |
| 3 | 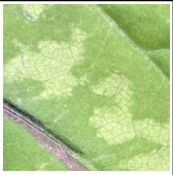 | Small portions of cell death are observed within the infiltrated area |
| 4 | 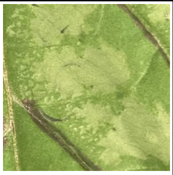 | Large patches of cell death is observed within the infiltrated area |
| 5 | 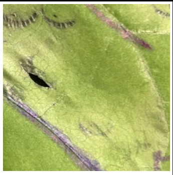 | Over half of the infiltrated area shows cell death |
| 6 | 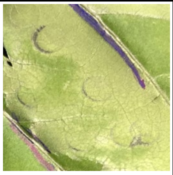 | Cell death is mostly observed in 80-90 % of the infiltrated area |
| 7 | 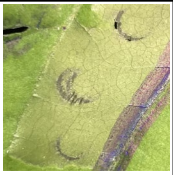 | Cell death is observed throughout the entire infiltrated area |

HR scoring criteria and representative photographs of HR in *S. americanum*. HR range is scored from 0 (no HR) to 7 (full HR). The scoring scales were revised based on the previous study [S2]

### Figure S3. Generation and confirmation of effector knockout, Related to Figure 5

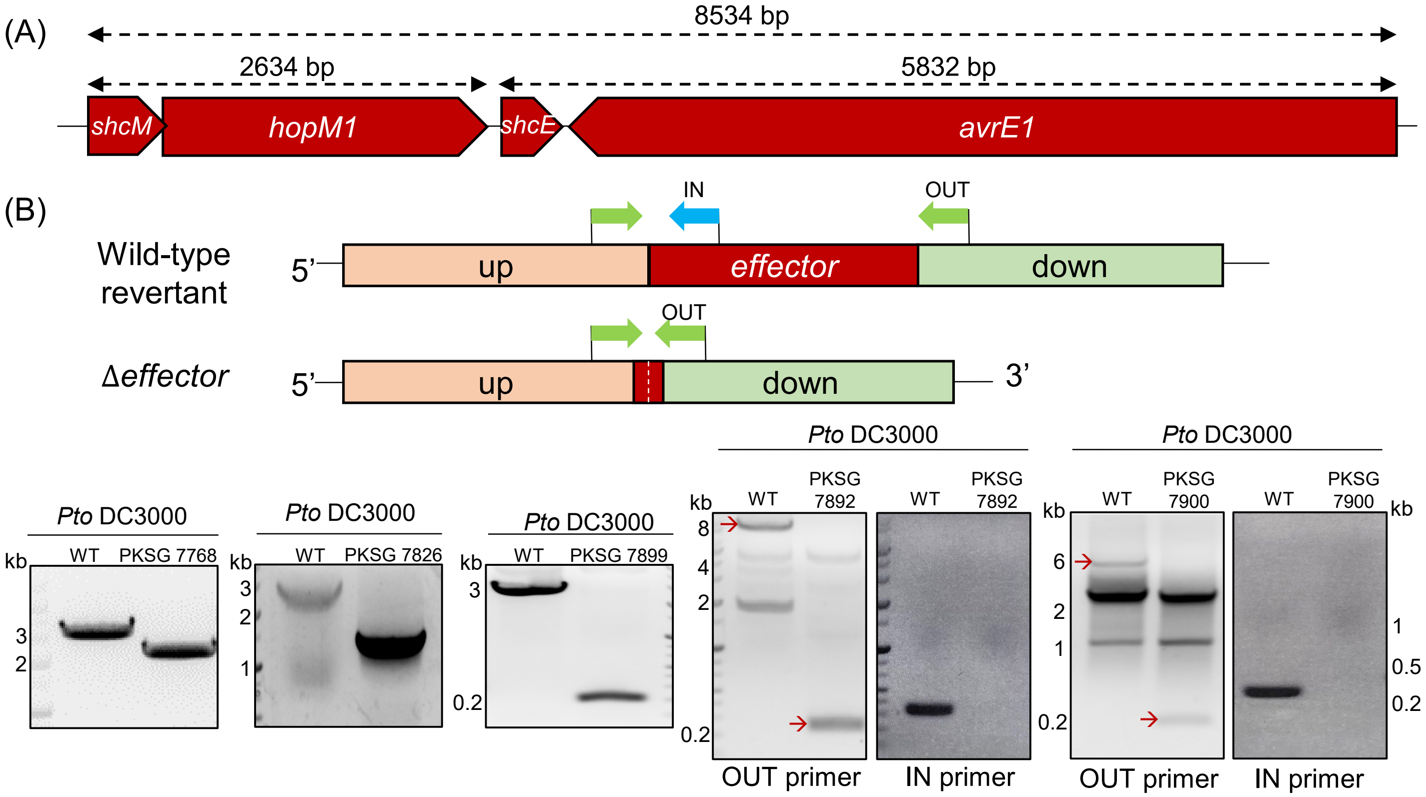


(C)

| | | Primer (PKSP) | Expected band size (bp) | |
|---|---|---|---|---|
| | | | WT | Δ*effector* |
| *hopC1* | OUT | 5987+5926 | 3126 | 2328 |
| *hopAA1-1* | OUT | 5983+6039 | 2864 | 1415 |
| *shcM-hopM1* | OUT | 6083+6093 | 2817 | 199 |
| *shcE-avrE1* | OUT | 6892+6056 | 5955 | 175 |
| | IN | 6892+6896 | 337 | no band |
| *shcM-hopM1:shcE-avrE1* | OUT | 6892+6093 | 8710 | 188 |
| | IN | 6892+6896 | 337 | no band |

(A) Schematic representation of effector knockout in *Pto* DC3000. For *hopC1* knockout, the upstream of *hopC1* flanking region was designed to be 1476 bp and the downstream was 1482 bp, with both regions containing 4 bp of *hopC1* gene 5' and 3' end. For *hopAA1-1* knockout, the upstream flanking region was 1201 bp and the downstream region was 1029 bp with 4 bp of the *hopAA1-1* gene 5' and 3' end, respectively. For *hopM1* knockout, the chaperone for HopM1 translocation (Badel *et al.*, 2003), *shcM* was deleted along with *hopM1*. The upstream of *shcM* flanking region was 1416 bp and the downstream was 1467 bp, each containing 4 bp of *shcM* 5' end and 4 bp of *hopM1* 3' end region. For *avrE1* knockout, both *avrE1* and its chaperone *shcE* were deleted together. The upstream flanking region was 1800 bp and the downstream flanking region was 1516 bp, each containing 4 bp of *shcE* 3' end and 3' end of *avrE1* genes, respectively. The upstream and the downstream flanking region constructs are cloned using Goldengate cloning into pK18mobsacB-GG compatible vector [S1]. A detailed effector gene knockout method is shown in the method section. (B) PCR confirmation of effector knockout in *Pto* DC3000. Primers for PCR were designed to be located in the flanking regions of the effector genes and the specific regions inside the effector genes. For OUT primers, the reduced band size indicates the deletion of the gene. For IN primers, the wild-type revertant shows a band, while the effector-deleted mutant does not. Green arrows indicate the OUT primers and the blue arrow indicates the IN primer used to confirm the knockout. The red arrows show the expected band size in the confirmation PCR. (C) The table shows the primer names and expected band sizes for the effector knockout. PKSP means the used primer name. The primer sequences are provided in Table S11.

Table S1. *Solanum americanum* accessions used in this study

| Working name (accession) | Species | HR triggered by *Pto* DC3000 |
|---|---|---|
| SP2273 | *S. americanum* | + |
| SP2275 | *S. americanum* | + |
| SP2297 | *S. americanum* | + |
| SP2298 | *S. americanum* | + |
| SP2299 | *S. americanum* | + |
| SP2300 | *S. americanum* | + |
| SP2302 | *S. americanum* | + |
| SP2304 | *S. americanum* | + |
| SP2305 | *S. americanum* | + |
| SP2306 | *S. americanum* | + |
| SP2308 | *S. americanum* | + |
| SP2359 | *S. americanum* | + |
| SP2360 | *S. americanum* | + |
| SP2361 | *S. americanum* | + |
| SP3050 | *S. americanum* | + |
| SP3398 | *S. americanum* | + |
| SP3405 | *S. americanum* | + |
| SP1123 | *S. americanum* | + |
| SP2268 | *S. americanum* | + |
| SP2269 | *S. americanum* | + |
| SP3051 | *S. americanum* | + |
| SP3389 | *S. americanum* | + |
| SP3399 | *S. americanum* | + |
| SP3401 | *S. americanum* | + |
| SP3402 | *S. americanum* | + |
| SP3403 | *S. americanum* | + |
| SP3404 | *S. americanum* | + |
| SP3406 | *S. americanum* | + |

'+' indicates HR development at 1day post infection. *Pto* DC3000 wild-type was infiltrated in a high concentration ($OD_{600nm}$=0.1) into *S. americanum* leaves using a needleless syringe.

### Table S2. Bacterial strains used in this study

| Bacterial strains | Relevant characteristics | Reference |
|---|---|---|
| *E.coli* DH5α | Wild-type; For cloning plasmids | Lab collection |
| *E.coli* HB101 | pRK2013; $Kan^R$ | [S3] |
| *A. tumafaciens* AGL1 | Wild-type; $Carb^R$ | Lab collection |
| *Pto* DC3000 | Wild-type; $Rif^R$ | [S4] |
| *Pto* DC3000 D36E | *Pto* DC3000 polymutant lacking 36 effectors; $Rif^R$, $Spec^R$ | [S5] |
| *Pto* DC3000 D29E | *Pto* DC3000 polymutant lacking 29 effectors; $Rif^R$, $Spec^R$ | [S5] |
| *Pto* DC3000 D18E | *Pto* DC3000 polymutant lacking 18 effectors; $Rif^R$ | [S6] |
| PKSG 4673 | Unmarked deletion of avrPto in *Pto* DC3000 using pK18mobsacB-GG-*avrPto*; *Pto* DC3000 Δ*avrPto*; $Rif^R$ | This study |
| PKSG 7065 | Unmarked deletion of *hopAM1-1* in PKSG 4673; *Pto* DC3000 Δ*avrPto*Δ*hopAM1-1*;$Rif^R$ | This study |
| PKSG 7903 | Unmarked deletion of *hopAM1-2* in PKSG 7065; *Pto* DC3000 Δ*avrPto*Δ*hopAM1-1*Δ*hopAM1-2*;$Rif^R$ | This study |
| PKSG 7377 | Unmarked deletion of *hopAD1* in PKSG 7903; *Pto* DC3000 Δ*avrPto*Δ*hopAM1-1*Δ*hopAM1-2*Δ*hopAD1*; $Rif^R$ | This study |
| PKSG 7768 | Unmarked deletion of hopC1 in PKSG 7377; *Pto* DC3000 Δ*avrPto*Δ*hopAM1-1*ΔhopAM1-2Δ*hopAD1*Δ*hopC1*; $Rif^R$ | This study |
| PKSG 7826 | Unmarked deletion of *hopAA1-1* in PKSG 7768; *Pto* DC3000 Δ*avrPto*Δ*hopAM1-1*Δ*hopAM1-2*Δ*hopAD1*Δ*hopC1*Δ*hopAA1-1*; $Rif^R$ | This study |
| PKSG 7899 | Unmarked deletion of *shcM : hopM1* in PKSG 7826; *Pto* DC3000 Δ*avrPto*Δ*hopAM1-1*Δ*hopAM1-2*Δ*hopAD1*Δ*hopC1*Δ*hopAA1-1* Δ*hopM1(shcM)*; $Rif^R$ | This study |
| PKSG 7900 | Unmarked deletion of *shcE : avrE1* in PKSG 7826; *Pto* DC3000 Δ*avrPto*Δ*hopAM1-1*Δ*hopAM1-2*Δ*hopAD1*Δ*hopC1*Δ*hopAA1-1* Δ*avrE1(shcE)*; $Rif^R$ | This study |
| PKSG 7892 | Unmarked deletion of *shcM : hopM1 : shcE : avrE1* in PKSG 7826; Δ*avrPto*Δ*hopAM1-1*Δ*hopAM1-2*Δ*hopAD1*Δ*hopC1*Δ*hopAA1-1* Δ*hopM1(shcM)*Δ*avrE1(shcE)*; $Rif^R$ | This study |

### Table S3. Plasmids used in this study

| Plasmids | Relevant characteristics | Reference |
|---|---|---|
| pICH41021 | *BsaI* restriction site mutagenized pUC19 vector | Lab collection |
| pICH86988 | Binary vector for transient gene expression consisting CaMV 35S promoter; $Kan^R$ | [S5] |
| pK18mobsacB-GG | Goldengate compatible pk18mobsacB vector; Suicide vector for effector knockout; $Kan^R$ | [S1], [S8] |
| pBBR1MCS-5B 662 | Goldengate compatible broad host range vector for effector delivery; $Gen^R$ | [S9] |
| pICH86988: *hopA1*: HA | *hopA1*$_{Pto\ DC3000}$: HA in pICH86988; $Kan^R$ | This study |
| pICH86988: *hopB1*: HA | *hopB1*$_{Pto\ DC3000}$: HA in pICH86988; $Kan^R$ | This study |
| pICH86988 *hopE1*: HA | *hopE1*$_{Pto\ DC3000}$: HA in pICH86988; $Kan^R$ | This study |
| pICH86988: *hopI1*: HA | *hopI1*$_{Pto\ DC3000}$: HA in pICH86988; $Kan^R$ | This study |
| pICH86988: *hopK1*: HA | *hopK1*$_{Pto\ DC3000}$: HA in pICH86988; $Kan^R$ | This study |
| pICH86988: *hopY1*: HA | *hopY1*$_{Pto\ DC3000}$: HA in pICH86988; $Kan^R$ | This study |
| pICH86988: *avrPto*: HA | *avrPto*$_{Pto\ DC3000}$: HA cloned in pICH86988; $Kan^R$ | This study |
| pICH86988: *avrPtoB*: HA | *avrPtoB*$_{Pto\ DC3000}$: HA cloned in pICH86988; $Kan^R$ | This study |
| pICH86988: *hopAD1*: HA | *hopAD1*$_{Pto\ DC3000}$: HA cloned in pICH86988; $Kan^R$ | This study |
| pICH86988: *hopAF1*: HA | *hopAF1*$_{Pto\ DC3000}$: HA cloned in pICH86988; $Kan^R$ | This study |
| pICH86988: *hopAM1-1*: HA | *hopAM1-1*$_{Pto\ DC3000}$: HA cloned in pICH86988; $Kan^R$ | This study |
| pICH86988: *hopU1*: HA | *hopU1* $_{Pto\ DC3000}$: HA cloned in pICH86988; $Kan^R$ | This study |
| pICH86988: *hopF2*: HA | *hopF2*$_{Pto\ DC3000}$: HA cloned in pICH86988; $Kan^R$ | This study |
| pICH86988: *hopH1*: HA | *hopH1* $_{Pto\ DC3000}$ : HA cloned in pICH86988; $Kan^R$ | This study |
| pICH86988: *hopC1*: HA | *hopC1*$_{Pto\ DC3000}$: HA cloned in pICH86988; $Kan^R$ | This study |
| pICH86988: *hopD1*: HA | *hopD1*$_{Pto\ DC3000}$: HA cloned in pICH86988; $Kan^R$ | This study |
| pICH86988: *hopQ1-1*: HA | *hopQ1-1*$_{Pto\ DC3000}$: HA cloned in pICH86988; $Kan^R$ | This study |
| pICH86988: *hopR1*: HA | *hopR1*$_{Pto\ DC3000}$: HA cloned in pICH86988; $Kan^R$ | This study |
| pICH86988: *hopN1*: HA | *hopN1*$_{Pto\ DC3000}$: HA cloned in pICH86988; $Kan^R$ | This study |
| pICH86988: *hopAA1-1*: HA | *hopAA1-1*$_{Pto\ DC3000}$: HA cloned in pICH86988; $Kan^R$ | This study |
| pICH86988: *hopM1*: HA | *hopM1*$_{Pto\ DC3000}$: HA cloned in pICH86988; $Kan^R$ | This study |
| pICH86988: *avrE1*: HA | *avrE1*$_{Pto\ DC3000}$: HA cloned in pICH86988; $Kan^R$ | This study |
| pICH86988: *hopAA1-2*: HA | *hopAA1-2*$_{Pto\ DC3000}$: HA cloned in pICH86988; $Kan^R$ | This study |
| pICH86988: *hopV1*: HA | *hopV1*$_{Pto\ DC3000}$: HA cloned in pICH86988; $Kan^R$ | This study |
| pICH86988: *hopAO1*: HA | *hopAO1*$_{Pto\ DC3000}$: HA cloned in pICH86988; $Kan^R$ | This study |
| pICH86988: *hopG1*: HA | *hopG1*$_{Pto\ DC3000}$: HA cloned in pICH86988; $Kan^R$ | This study |
| pICH86988: *hopX1*: HA | *hopX1*$_{Pto\ DC3000}$: HA cloned in pICH86988; $Kan^R$ | This study |
| pICH86988: *hopO1-1*: HA | *hopO1-1*$_{Pto\ DC3000}$: HA cloned in pICH86988; $Kan^R$ | This study |
| pICH86988: *hopT1-1*: HA | *hopT1-1*$_{Pto\ DC3000}$: HA cloned in pICH86988; $Kan^R$ | This study |
| pk18mobsacB-GG - *avrPto* | *avrPto* up: down flanking regions cloned in pk18mobsacB-GG; $Kan^R$ | This study |
| pk18mobsacB-GG - *hopAM1-1* | *hopAM1-1* up: down flanking regions cloned in pk18mobsacB-GG; $Kan^R$ | This study |
| pk18mobsacB-GG - *hopAM1-2* | *hopAM1-2* up: down flanking regions cloned in pk18mobsacB-GG, $Kan^R$ | This study |
| pk18mobsacB-GG - *hopAD1* | *hopAD1* up: down flanking regions cloned in pk18mobsacB-GG; $Kan^R$ | This study |
| pk18mobsacB-GG - *hopC1* | *hopC1* up: down flanking regions cloned in pk18mobsacB-GG; $Kan^R$ | This study |
| pk18mobsacB-GG - *hopAA1-1* | *hopAA1-1* up: down flanking regions cloned in pk18mobsacB-GG, $Kan^R$ | This study |
| pk18mobsacB-GG - *shcM : hopM1* | *shcM/hopM1* up: down flanking regions cloned in pk18mobsacB-GG; $Kan^R$ | This study |
| pk18mobsacB-GG - *shcE : avrE1* | *shcE/avrE1* up: down flanking regions cloned in pk18mobsacB-GG; $Kan^R$ | This study |
| pk18mobsacB-GG - *shcM* : *hopM1*: *shcE : avrE1* | *shcM/avrE1* up: down flanking regions cloned in pk18mobsacB-GG; $Kan^R$ | This study |
| pBBR1MCS-5B 662 - *avrPto* promoter *: shcE : avrE1* : HA | *avrPto* promoter, *shcE, avrE1*: HA cloned in pBBR1MCS 5B:662; $Gen^R$ | This study |

Table S4. Raw data of *in planta* bacterial growth, Related to Figure 1

| | Replicate 1 | | | Replicate 2 | | | Replicate 3 | | | |
|---|---|---|---|---|---|---|---|---|---|---|
| | Repeat No. | | | Repeat No. | | | Repeat No. | | | |
| | 1 | 2 | 3 | 1 | 2 | 3 | 1 | 2 | 3 | 4 |
| *Pto* D36E | 4.176 | 4.477 | 4.000 | 3.699 | 4.146 | 3.000 | | | | |
| *Pto* D29E | 2.699 | 4.556 | 4.000 | 3.000 | 3.301 | 4.301 | 4.000 | 3.176 | 4.204 | 3.301 |
| *Pto* D18E | 4.618 | 4.618 | 4.423 | 4.301 | 4.477 | 4.398 | 4.699 | 4.602 | 4.477 | 4.544 |
| *Pto* DC3000 Wild-type | 5.332 | 5.041 | 4.778 | 4.398 | 4.699 | 4.929 | 5.290 | 4.653 | 4.954 | 4.653 |

Table S5. Type III effector expected protein size, Related to Figure 2 and Figure 4

| | Effector | Size (kDa) |
|---|---|---|
| 1 | HopA1 | 49 |
| 2 | HopAD1 | 76 |
| 3 | HopAF1 | 38 |
| 4 | HopAM1 | 38 |
| 5 | HopB1 | 57 |
| 6 | HopE1 | 31 |
| 7 | AvrPto | 25 |
| 8 | AvrPtoB | 66 |
| 9 | HopI1 | 60 |
| 10 | HopK1 | 44 |
| 11 | HopY1 | 38 |
| 12 | HopAA1-1 | 61 |
| 13 | HopAA1-2 | 58 |
| 14 | HopAO1 | 55 |
| 15 | AvrE1 | 202 |
| 16 | HopC1 | 37 |
| 17 | HopD1 | 82 |
| 18 | HopR1 | 217 |
| 19 | HopG1 | 62 |
| 20 | HopH1 | 31 |
| 21 | HopM1 | 82 |
| 22 | HopN1 | 46 |
| 23 | HopO1-1 | 38 |
| 24 | HopQ1-1 | 56 |
| 25 | HopF2 | 29 |
| 26 | HopT1-1 | 48 |
| 27 | HopU1 | 35 |
| 28 | HopV1 | 50 |
| 29 | HopX1 | 51 |

### Table S6. Raw data of *in planta* bacterial growth, Related to Figure 3

| | Replicate 1 $Log_{10}(CFU/cm^2)$ | | | Replicate 2 $Log_{10}(CFU/cm^2)$ | | | | Replicate 3 $Log_{10}(CFU/cm^2)$ | | | Replicate 4 $Log_{10}(CFU/cm^2)$ | | | | Replicate 5 $Log_{10}(CFU/cm^2)$ | | | |
|---|---|---|---|---|---|---|---|---|---|---|---|---|---|---|---|---|---|---|
| | Repeat No. | | | Repeat No. | | | | Repeat No. | | | Repeat No. | | | | Repeat No. | | | |
| | 1 | 2 | 3 | 1 | 2 | 3 | 4 | 1 | 2 | 3 | 1 | 2 | 3 | 4 | 1 | 2 | 3 | 4 |
| *Pto* D36E | 3.929 | 3.544 | 3.653 | 3.653 | 3.544 | 3.740 | 2.699 | 4.079 | 3.978 | 2.699 | | | | | | | | |
| *Pto* DC3000 Wild-type | 5.380 | 5.204 | 4.653 | 5.041 | 5.312 | 5.190 | 5.061 | 5.190 | 5.190 | 5.041 | 4.778 | 4.000 | 4.000 | 4.740 | 4.929 | 4.000 | 3.699 | 4.398 |
| PKSG 4673 | 5.978 | 6.114 | 5.929 | | | | | 5.875 | 5.778 | 5.778 | 6.301 | 5.477 | 5.778 | 6.190 | 6.000 | 6.602 | 6.000 | 6.301 |
| PKSG 7065 | 6.778 | 6.398 | 6.602 | 6.398 | 6.176 | 6.041 | 5.602 | 6.279 | 5.845 | 6.021 | 6.699 | 6.903 | 6.301 | 6.000 | 6.778 | 5.699 | 6.000 | 6.000 |
| PKSG 7903 | | | | | | | | 6.000 | 6.130 | 5.813 | 6.000 | 6.740 | 6.176 | 6.398 | 6.954 | 7.000 | 5.699 | 6.544 |
| PKSG 7377 | | | | 6.407 | 6.447 | 6.079 | 5.544 | 6.243 | 6.217 | 5.813 | 5.699 | 6.477 | 5.699 | 6.477 | 6.371 | 5.845 | 5.602 | 6.021 |

### Table S7. Raw data of hypersensitive response scoring, Related to Figure 3

| | Replicate 1 | | | | Replicate 2 | | | | Replicate 3 | | | Replicate 4 | | Replicate 5 | | | | Replicate 6 | | | | Replicate 7 | | | | |
|---|---|---|---|---|---|---|---|---|---|---|---|---|---|---|---|---|---|---|---|---|---|---|---|---|---|---|
| | Repeat No. | | | | Repeat No. | | | | Repeat No. | | | Repeat No. | | Repeat No. | | | | Repeat No. | | | | Repeat No. | | | | |
| | 1 | 2 | 3 | 4 | 1 | 2 | 3 | 4 | 1 | 2 | 3 | 1 | 2 | 1 | 2 | 3 | 4 | 1 | 2 | 3 | 4 | 1 | 2 | 3 | 4 | 5 |
| *Pto* D36E | 0 | 0 | 0 | 0 | 0 | 0 | 2 | 3 | 0 | 0 | 0 | 0 | 0 | 0 | 0 | 0 | 0 | 0 | 0 | 0 | 0 | 0 | 0 | 0 | 0 | 0 |
| *Pto* DC3000 Wild-type | 7 | 7 | 7 | 7 | 7 | 7 | 7 | 7 | 7 | 7 | 7 | 7 | 7 | 7 | 7 | 7 | 7 | 7 | 7 | 7 | 7 | 7 | 7 | 7 | 7 | 7 |
| PKSG 4673 | 7 | 7 | 7 | 7 | 7 | 7 | 7 | 7 | 7 | 7 | 7 | / | / | 7 | 7 | 6 | 6 | 7 | 6 | 6 | 2 | 6 | 6 | 6 | 7 | 7 |
| PKSG 7065 | / | / | / | / | / | / | / | / | 7 | 7 | 7 | 7 | 7 | 7 | 7 | 7 | 7 | 7 | 7 | 1 | 5 | 7 | 7 | 7 | 7 | 7 |
| PKSG 7903 | / | / | / | / | / | / | / | / | / | / | / | 7 | 7 | 7 | 7 | 7 | 6 | 6 | 6 | 1 | 1 | 6 | 6 | 7 | 7 | 7 |
| PKSG 7377 | / | / | / | / | / | / | / | / | / | / | / | / | / | 7 | 7 | 7 | 7 | 7 | 5 | 1 | 1 | 7 | 7 | 7 | 7 | 7 |

### Table S8. Raw data of *in planta* bacterial growth, Related to Figure 5

| | Replicate 1 Log$_{10}$(CFU/cm$^2$) | | | | Replicate 2 Log$_{10}$(CFU/cm$^2$) | | | | Replicate 3 Log$_{10}$(CFU/cm$^2$) | | | Replicate 4 Log$_{10}$(CFU/cm$^2$) | | | | Replicate 5 Log$_{10}$(CFU/cm$^2$) | | | Replicate 6 Log$_{10}$(CFU/cm$^2$) | | |
|---|---|---|---|---|---|---|---|---|---|---|---|---|---|---|---|---|---|---|---|---|---|
| | Repeat No. | | | | Repeat No. | | | | Repeat No. | | | Repeat No. | | | | Repeat No. | | | Repeat No. | | |
| | 1 | 2 | 3 | 4 | 1 | 2 | 3 | 4 | 1 | 2 | 3 | 1 | 2 | 3 | 4 | 1 | 2 | 3 | 1 | 2 | 3 |
| *Pto* D36E | 3.778 | 3.602 | 2.079 | 2.784 | | | | | 3.929 | 4.021 | 3.778 | | | | | 3.544 | 3.000 | 3.398 | 3.544 | 3.000 | 3.398 |
| *Pto* DC3000 Wild-type | 4.699 | 4.176 | 5.230 | 5.146 | 4.602 | 4.398 | 4.544 | 4.778 | 5.161 | 4.903 | 5.021 | 4.740 | 5.146 | 4.845 | 4.301 | 4.845 | 4.929 | 5.161 | 4.845 | 4.929 | 5.161 |
| PKSG 7377 | 6.130 | 6.439 | 6.176 | 6.190 | | | | | 5.000 | 6.000 | 5.740 | 5.653 | 5.398 | 5.602 | 5.176 | 6.176 | 6.301 | 6.544 | 6.176 | 6.301 | 6.544 |
| PKSG 7768 | | | | | 6.130 | 6.079 | 5.954 | 6.322 | 6.491 | 6.484 | 6.230 | 5.740 | 5.903 | 5.602 | 6.146 | 5.653 | 5.301 | 6.061 | 5.653 | 5.301 | 6.061 |
| PKSG 7826 | | | | | 6.130 | 6.243 | 6.176 | 6.061 | 6.778 | 6.929 | 6.778 | 6.021 | 5.544 | 5.929 | 6.322 | 6.301 | 6.813 | 6.176 | 6.301 | 6.813 | 6.176 |
| PKSG 7899 | | | | | | | | | 5.114 | 4.176 | 5.362 | 5.778 | 5.845 | 5.978 | 5.954 | 6.176 | 6.477 | 6.000 | 6.176 | 6.477 | 6.000 |
| PKSG 7900 | | | | | | | | | 5.279 | 5.000 | 4.477 | 4.699 | 5.000 | 5.544 | 5.398 | 6.021 | 5.875 | 5.653 | 6.021 | 5.875 | 5.653 |
| PKSG 7892 | 4.301 | 4.176 | 4.544 | 4.699 | | | | | 5.371 | 5.114 | 4.477 | 5.978 | 5.000 | 5.301 | 4.699 | 4.312 | 3.699 | 4.415 | 4.312 | 3.699 | 4.415 |

### Table S9. Raw data of hypersensitive response scoring, Related to Figure 5

| | Replicate 1 | | | | Replicate 2 | | | | Replicate 3 | | | | Replicate 4 | | | | | Replicate 5 | | | |
|---|---|---|---|---|---|---|---|---|---|---|---|---|---|---|---|---|---|---|---|---|---|
| | Repeat No. | | | | Repeat No. | | | | Repeat No. | | | | Repeat No. | | | | | Repeat No. | | | |
| | 1 | 2 | 3 | 4 | 1 | 2 | 3 | 4 | 1 | 2 | 3 | 4 | 1 | 2 | 3 | 4 | 5 | 1 | 2 | 3 | 4 |
| *Pto* D36E | 0 | 0 | 0 | 0 | 0 | 0 | 0 | 0 | 0 | 0 | 0 | 0 | 0 | 0 | 0 | 0 | 0 | 0 | 0 | 0 | 0 |
| *Pto* DC3000 Wild-type | 7 | 7 | 7 | 7 | 7 | 7 | 7 | 7 | 7 | 7 | 7 | 7 | 6 | 7 | 7 | 7 | 7 | 7 | 7 | 7 | 7 |
| PKSG 7377 | | | | | | | | | 7 | 6 | 7 | 6 | 1 | 1 | 6 | 6 | 6 | 7 | 7 | 7 | 7 |
| PKSG 7768 | | | | | | | | | 6 | 6 | 4 | 3 | 1 | 2 | 2 | 3 | 3 | 4 | 4 | 4 | 4 |
| PKSG 7826 | | | | | | | | | 2 | 4 | 3 | 2 | 3 | 3 | 3 | 3 | 3 | 1 | 4 | 4 | 6 |
| PKSG 7899 | | | | | | | | | 6 | 3 | 3 | 2 | 0 | 3 | 3 | 3 | 4 | 4 | 4 | 4 | 6 |
| PKSG 7900 | | | | | 2 | 3 | 6 | 6 | 2 | 0 | 1 | 0 | 0 | 2 | 2 | 2 | 3 | 3 | 3 | 3 | 5 |
| PKSG 7892 | 0 | 0 | 2 | 0 | 2 | 3 | 4 | 6 | 2 | 0 | 0 | 0 | 0 | 0 | 0 | 0 | 0 | 0 | 0 | 0 | 1 |

Table S10. number of species or pathovars analyzed in this study, Related to Figure 7

| Taxon | number of species or pathovar analyzed |
|---|---|
| *Pseudomonas viridiflava* | 10 |
| *Pseudomonas tremae* | 2 |
| *Pseudomonas syringae* | 23 |
| *Pseudomonas syringae* pv. *tomato* | 6 |
| *Pseudomonas syringae* pv. *tagetis* | 1 |
| *Pseudomonas syringae* pv. *syringae* | 10 |
| *Pseudomonas syringae* pv. *pisi* | 1 |
| *Pseudomonas syringae* pv. *maculicola* | 2 |
| *Pseudomonas syringae* pv. *lapsa* | 1 |
| *Pseudomonas syringae* pv. *helianthi* | 1 |
| *Pseudomonas syringae* pv. *cerasicola* | 1 |
| *Pseudomonas syringae* pv. *avii* | 1 |
| *Pseudomonas syringae* pv. *atrofaciens* | 2 |
| *Pseudomonas syringae* pv. *antirrhini* | 1 |
| *Pseudomonas syringae* pv. *actinidifoliorum* | 1 |
| *Pseudomonas syringae* pv. *actinidiae* | 18 |
| *Pseudomonas syirngae group genomosp. 3* | 1 |
| *Pseudomonas savastanoi* | 5 |
| *Pseudomonas lijiangensis* | 1 |
| *Pseudomonas fuscovaginae* | 2 |
| *Pseudomonas ficuserectae* | 1 |
| *Pseudomonas coronafaciens* | 3 |
| *Pseudomonas cichorii* | 2 |
| *Pseudomonas cannabina* | 2 |
| *Pseudomonas avellanae* | 3 |
| *Pseudomonas asturiensis* | 1 |
| *Pseudomonas amygdali* | 15 |
| **Total strains** | **117** |

### Supplemental references


[S1] Jayaraman, J., Yoon, M., Applegate, E.R., Stroud, E.A., and Templeton, M.D. (2020). AvrE1 and HopR1 from Pseudomonas syringae pv. actinidiae are additively required for full virulence on kiwifruit. Molecular Plant Pathology 21:1467-1480. https://doi.org/10.1111/mpp.12989.

[S2] Ahn, Y.J., Kim, H., Choi, S., Mazo-Molina, C., Prokchorchik, M., Zhang, N., Kim, B., Mang, H., Koehler, N., and Kim, J. (2023). Ptr1 and ZAR1 immune receptors confer overlapping and distinct bacterial pathogen effector specificities. New Phytologist 239:1935-1953.
[S3] Figurski, D.H., and Helinski, D.R. (1979). Replication of an origin-containing derivative of plasmid RK2 dependent on a plasmid function provided in trans. Proc Natl Acad Sci U S A 76:1648-1652. 10.1073/pnas.76.4.1648.

[S4] Cuppels, D.A. (1986). Generation and characterization of Tn 5 insertion mutations in Pseudomonas syringae pv. tomato. Applied and environmental microbiology 51:323-327

[S5] Wei, H.-L., Chakravarthy, S., Mathieu, J., Helmann, T.C., Stodghill, P., Swingle, B., Martin, G.B., and Collmer, A. (2015). Pseudomonas syringae pv. tomato DC3000 type III secretion effector polymutants reveal an interplay between HopAD1 and AvrPtoB. Cell host & microbe 17:752-762.

[S6] Kvitko, B.H., Park, D.H., Velásquez, A.C., Wei, C.-F., Russell, A.B., Martin, G.B., Schneider, D.J., and Collmer, A. (2009). Deletions in the repertoire of Pseudomonas syringae pv. tomato DC3000 type III secretion effector genes reveal functional overlap among effectors. PLoS pathogens 5:e1000388.

[S7] Weber, E., Engler, C., Gruetzner, R., Werner, S., and Marillonnet, S. (2011). A modular cloning system for standardized assembly of multigene constructs. PloS one 6:e16765

[S8] Schäfer, A., Tauch, A., Jäger, W., Kalinowski, J., Thierbach, G., and Pühler, A. (1994). Small mobilizable multi-purpose cloning vectors derived from the Escherichia coli plasmids pK18 and pK19: selection of defined deletions in the chromosome of Corynebacterium glutamicum. Gene 145:69-73. 10.1016/0378-1119(94)90324-7.

[S9] Jayaraman, J., Choi, S., Prokchorchik, M., Choi, D.S., Spiandore, A., Rikkerink, E.H., Templeton, M.D., Segonzac, C., and Sohn, K.H. (2017). A bacterial acetyltransferase triggers immunity in Arabidopsis thaliana independent of hypersensitive response. Scientific Reports 7:3557.
